# Leveraging spatio-temporal genomic breeding value estimates of dry matter yield and herbage quality in ryegrass via random regression models

**DOI:** 10.1101/2022.05.01.489357

**Authors:** Elesandro Bornhofen, Dario Fè, Ingo Lenk, Morten Greve, Thomas Didion, Christian Sig Jensen, Torben Asp, Luc Janss

## Abstract

Joint modeling of correlated multi-environment and multi-harvest data of perennial crop species may offer advantages in prediction schemes and a better understanding of the underlying dynamics in space and time. The goal of the present study was to investigate the relevance of incorporating the longitudinal dimension of within-season multiple harvests of biomass yield and nutritive quality traits of forage perennial ryegrass (*Lolium perenne* L.) in a reaction norm model setup that additionally accounts for genotype-environment interactions. Genetic parameters and accuracy of genomic breeding value predictions were investigated by fitting three random regression (random coefficients) linear mixed models (gRRM) using Legendre polynomial functions to the data. All models accounted for heterogeneous residual variance and moving average-based spatial adjustments within environments. The plant material consisted of 381 bi-parental family pools and four check varieties of diploid perennial ryegrass evaluated in eight environments for biomass yield and nutritive quality traits. The longitudinal dimension of the data arose from multiple harvests performed four times annually. The specified design generated a total of 16,384 phenotypic data points for each trait. Genomic DNA sequencing was performed using DNA nanoball-based technology (DNBseq) and yielded 56,645 single nucleotide polymorphisms (SNPs) which were used to calculate the allele frequency-based genomic relationship matrix used in all genomic random regression models. Biomass yield’s estimated additive genetic variance and heritability values were higher in later harvests. The additive genetic correlations were moderate to low in early measurements and peaked at intermediates, with fairly stable values across the environmental gradient, except for the initial harvest data collection. This led to the conclusion that complex genotype-by-environment interaction (G×E) arises from spatial and temporal dimensions in the early season, with lower re-ranking trends thereafter. In general, modeling the temporal dimension with a second-order orthogonal polynomial in the reaction norm mixed model framework improved the accuracy of genomic estimated breeding value prediction for nutritive quality traits, but no gain in prediction accuracy was detected for dry matter yield. This study leverages the flexibility and usefulness of gRRM models for perennial ryegrass research and breeding and can be readily extended to other multi-harvest crops.

## Introduction

The largest portion of the world’s agricultural area is covered by grasses, with grazing systems spread over 22% of the Earth’s ice-free land surface (Ramankutty et al., 2008). Among perennial forage grasses, ryegrass (*Lolium perenne* L.) is the most widely sown species in temperate climate zones. Its breeding process, however, is not straightforward and can take 10–15 years from the concept to the release of a new cultivar (Lee et al., 2012), restricting genetic gains to levels below the ones estimated for other major crops (McDonagh et al., 2016). A first complication in Perennial ryegrass (2n = 2x = 14 [natural occurency] or 2n = 4x = 28 [induced]) breeding arises from being a cross-pollinated, self-incompatible species, that is bred in families or populations of related, but genetically heterogeneous, offspring, and with large effective population size (*Ne*). Molecular breeding can be deployed here aiming to overcome some of the constraints and ensure that resources will be allocated only to superior candidates. In spite of the challenges of implementing genomic selection in outbred species with large *Ne* as in the case of ryegrass (Lin et al., 2014), promising results have been shown for genome-wide marker-based prediction of dry matter yield (Guo et al., 2018; Arojju et al., 2020), heading date (Fè et al., 2015), and nutritive quality traits (Arojju et al., 2020).

A second complication arises from ryegrass being a perennial biomass crop, that can be harvested multiple times within a year, and across years. As is common in plant breeding, selection candidates will additionally be evaluated in multi-environment trials, and may show genotype-by-environment interactions (G×E) that can exhibit plasticity with developmental stage within a year and plant age across years. This underlying complex (co)variance structure from the multidimensional space involved is rarely explored in multiple harvest crop species, which results in a lack of knowledge about G×E effects at different crop ages across environments and vice versa. Sophisticated methods to account for data coming from multiple harvests (Faveri et al., 2015) and multiple environments (Malosetti et al., 2016) are available but studies combining these dimensions remain scarce. A few empirical studies show important results for the combined analysis of multiple harvest-location trials (Ferrão et al., 2017; Giri et al., 2019). In this context, variance and covariance modeling usually relies on stationary parametric correlation and factor-analytic structures. However, no study to date has investigated the covariance structure defined in random regression (RRM) models (Meyer & Kirkpatrick, 2005) to the analysis of multiple harvest-location data from perennial crops.

The standard approach to analyse multi-harvest data is to use the total annual biomass production as the response variable. This is rather a simplification and may miss biologically important aspects shaping genotypes’ response over time, and ignores time-dependent covariates. Throughout the production year, environmental variables change and so do the genotypes’ responses, given its level of plasticity. This linear or higher-order reaction norm can be expressed through a regression on functions of continuous covariables in the mixed model framework. Such models have been extensively used in animal breeding to model lactation curves given their longitudinal characteristics, and are referred to as test-day milk yield models. They are computationally efficient in describing trait trajectories as covariance functions (e.g., orthogonal polynomials and splines) explicitly defining the (co)variance between records using fewer parameters (Kirkpatrick et al., 1990; Meyer, 1998), and can be extended to multiple trait analysis. The advances and applications of random regression models were recently reviewed by (Oliveira et al., 2019) and include an overview of genomic prediction and genome-wide association studies based on genomic random regression models (gRRM). The use of gRRM in plants has been limited but is gaining traction due to the advent of high-throughput phenomics platforms that generate temporal phenotypes (Campbell et al., 2019; Momen et al., 2019; Moreira et al., 2021; Sun et al., 2017; Campbell et al., 2018) and to the availability of traits recorded over a continuous e.g., harvest time (do Amaral Santos de Carvalho Rocha et al., 2018) and environment (Ly et al., 2018; Marchal et al., 2019). These authors reported improvements when fitting gRRM models for quantitative genetic studies, demonstrating its efficacy for the analysis of complex and longitudinal traits. In addition, prediction accuracies of longitudinal traits were shown to be higher when simultaneously including an environmental-dependent covariate e.g., temperature-humidity index, in animal studies (Bohlouli et al., 2019). Therefore, gRRM models may be suited to accommodate the kind of data generated by multi-environment evaluation of ryegrass families.

Yearly growth and regrowth dry matter yield sum of ryegrass is the target breeding trait. However, there is a growing interest in the improvement of nutritive quality parameters e.g, dry matter digestibility (D-value), as it leads to increased animal performance (Miller et al., 2001). Forage quality traits are essentially time-dependent, showing substantial temporal variation as above-ground biomass is removed by sequential harvests or by animal grazing (McGrath et al., 2013), meaning they can be considered infinitely dimensional traits. The shape of their curves can be modeled by orthogonal polynomial functions, a way to model variances and covariances of a longitudinal trait (Schaeffer, 2016). Also, the existence of significant correlations among forage quality traits (Jafari et al., 2003; McGrath et al., 2013; Wang et al., 2015) implies that their trajectories are interrelated across seasonal plant growth and development. Therefore, random coefficient models using Legendre polynomials can be an attractive approach to describe the genetic variation in repeated measurements of ryegrass quality traits.

Since the conceptualization of the term reaction norm by Woltereck (1909) and the later well-known study from Finlay & Wilkinson (1963) describing phenotypes of a quantitative trait as a continuous function of a changing environment, substantial progress has been made towards the development of reaction norm models for characterization of G×E in animal and plant breeding (Su et al., 2006; Lian & de los Campos, 2016). Reaction norm models can outperform conventional models in terms of accuracy of breeding value estimates due to better exploitation of information from the target and adjacent environments (Mulder, 2016). Estimating population-level responses, individual-level reaction norms with varying coefficients, and the flexibility of model virtually any shape of linear or nonlinear response to a changing environment, makes gRRM models a powerful statistical tool to dissect plant plasticity (Arnold et al., 2019). In this study, reaction norm models were explored considering environment- and time-dependent covariates aiming to investigate the effect of modeling the temporal dimension on the relative accuracy of prediction of ryegrass dry matter yield and spectroscopy-based quality traits. In addition, discussions are elaborated regarding genetic parameters profiles, G×E interaction across sources of variation, and its relevance for ryegrass breeding.

## Material and Methods

### Plant material, field experiments, and phenotypes

To accomplish the goal of leveraging the usage of spatio-temporal data for genomic prediction in ryegrass, the data from the GreenSelect project was used. The dataset is characterized by 381 F_2_ family pools and four commercial checks of diploid perennial ryegrass (Toddington, Sputnik, Boyne, and Abosan 1), all developed by the breeding company DLF Seeds A/S. The F_2_ families were obtained by crossing 104 parents in two unconnected sparse diallel crossing schemes, instead of producing all *n* (*n−*1)*/*2 pairwise possible outcomes. The larger diallel (diallel A) comprises 88 parents, yielding 335 crosses, while the small one (diallel B) consisted of 16 parents and a total of 46 crosses (Figure 1, A). Parental plants were clonally propagated to allow the same genetic material to be used in more than one cross. Following one round of seed multiplication, two field experiments were carried out in two locations in Denmark (Figure 1, B). The experiments were sown during the fall season of 2018 at a seed rate of 22 kg ha^-1^. The first harvest event took place during the spring of 2019 when plants reached the boot (R0) growth stage (Moore et al., 1991). Subsequent harvests were performed in intervals of *∼* 5 weeks. Four harvests of fresh biomass were mechanically performed each year using a HALDRUP plot combine for a total of two testing years (2019 and 2020). Each field experiment layout consisted of entries assigned to one or two replicates within two nitrogen (N) availability conditions: i) normal N rate (380 kg^-1^ N ha^-1^ yr^-1^), and ii) low N rate (280 kg^-1^ N ha^-1^ yr^-1^). For the normal availability, 152 kg N ha^-1^ were applied in early spring at the beginning of growth, 114 kg N ha^-1^ after the first cut, 76 kg N ha^-1^ after the second, and 38 kg N ha^-1^ after the third cut. In the low-N condition, 112 kg N ha^-1^ were applied in early spring at the beginning of growth, 84 kg N ha^-1^ after the first cut, 56 kg N ha^-1^ after the second, and 28 kg N ha^-1^ after the third cut. Irrigation was supplied right after every N fertilization. Therefore, 1,024 experimental units (plots) of the size of 12.5 m^2^ were evaluated in location 1 in an irregular rectangular grid of 144 rows by eight columns, and the same number, but with a larger size (13.5 m^2^), in location 2 and laid out in a regular rectangular grid of 128 rows by eight columns.

**Figure 1:**
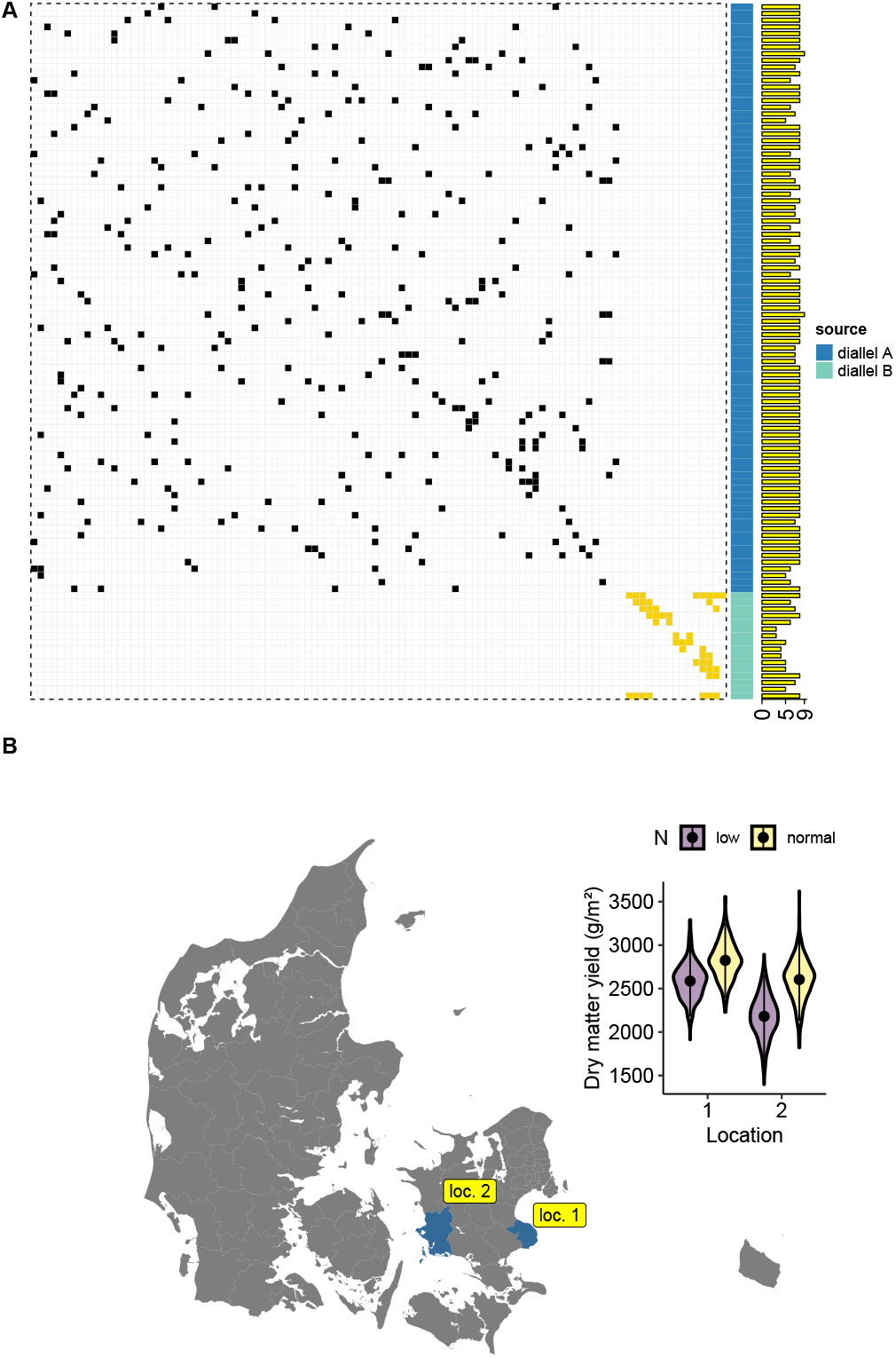
Graphical representation of two partial diallels crosses producing 381 F_2_ ryegrass families (A) and a map of the country of Denmark highlighting the two testing sites (B). Barplot in A shows the total number of realized crosses with each parental genotype. The map in B is annotated with a violin plot where black dots represent the overall mean for the cumulative two years dry matter yield at two nitrogen levels.

Phenotypes were recorded for eight traits at each harvest. Dry matter yield (DMY), expressed in grams of dry matter per squared meter, was assessed after correcting for moisture content. The seven nutritive quality traits are acid detergent fiber (ADF), acid deterged lignin (ADL), dry matter digestibility (DMDig), neutral deterged fiber (NDF), digestible NDF (NDFD), protein (Prot), and water-soluble carbohydrate (WSC); all measured as a percentage of DMY, except for NDFD, which is measured as proportion of NDF. Moisture and quality parameters were measured via a near-infrared spectrometer onboard the plot combine.

### Genomic data

All F_2_ families and check cultivars were subjected to the genotyping-by-sequence (GBS) process for genome-wide single nucleotide polymorphisms (SNPs) discovery. Plant genomic DNA was extracted from young leaves of bulks with several plants on a Tecan Freedom EVO robotic system (Tecan, Switzerland) using the MolGen PurePrep Leaf Kit (MolGen, The Netherlands), adapted for automated extraction. The DNA concentrations were normalized and digested with *Ape*KI (5-bp recognition site) and *Pst*I (6-bp recognition site) restriction enzymes during library preparation. The GBS libraries were prepared according to the protocol described by Poland et al. (2012) with minor modifications. Libraries were sequenced by Beijing Genomics Institute (BGI Tech Solutions, China) using DNA nanoball (DNBseq) high-throughput sequencing solution with SE100 reads. Sequence reads from 385 entries were trimmed to remove adaptors (using Scythe 0.994 - https://github.com/vsbuffalo/scythe) and low-quality reads (using Sickle 1.33 - https://github.com/najoshi/sickle), with the quality threshold for trimming equal to 20. Demultiplexing was performed using Sabre 1.00 (https://github.com/najoshi/sabre) after discarding reads with lengths inferior to 40 bp. Demultiplexed reads were mapped to the draft *Lolium perenne* sequence assembly (Byrne et al., 2015) using BWA 0.7.5 (Li & Durbin, 2010). Variant calling was performed using GATK 3.2.2 (McKenna et al., 2010) and only biallelic sites were retained for further analysis. The final set of SNPs was achieved after several tests of filtering parameters and inspections of PCA plots and consists of 56,645 markers. This set was reached following the exclusion of markers with a missing rate above 0.5, minor allele frequency lower than 0.05, mean read depth lower than one, and entries with more than 0.9 of missing data.

The similarity between every two entries was given by the genomic kernel of the form described in equation 1, which is a genomic relationship matrix (GMR) [method 1, VanRaden (2008)] modified to use allele frequencies as described by Ashraf et al. (2014).

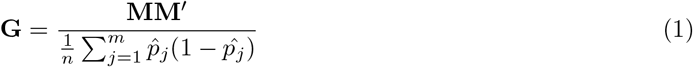

where **M** is a matrix of mean-centered allele frequencies and the cross-product of this matrix is further divided by a scaling parameter; *n* is the ploidy number of the breeding material and takes a value equal to four once diploid parents were crossed to generate the evaluated F_2_ families; *m* refers to the number of markers, and 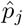 is the frequency of the *j*th marker. The diagonal of the relationship matrix **G** was then corrected by scaling down its elements according to an inflation parameter for each entry, which only depends on the ploidy number and the average S_T_ (coverage depth) of the sample (Cericola et al., 2018). A visual representation of the genomic relationship matrix is displayed as a heatmap in Figure 2 alongside annotations for trait means for every entry.

**Figure 2:**
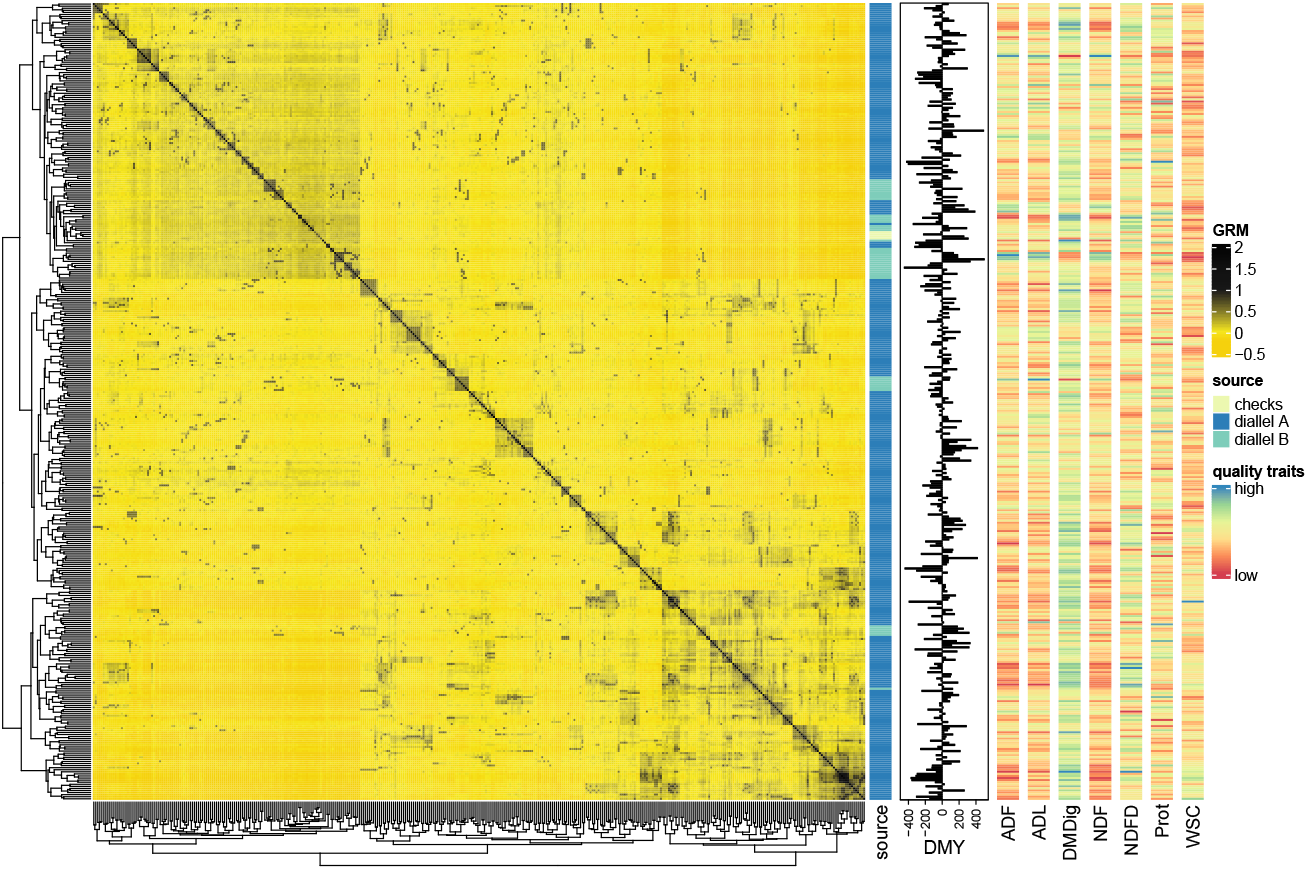
Heatmap of the genomic relationship matrix (GMR) annotated with family source information and phenotypic data from mean-centered dry matter yield (DMY) [barplot] and seven nutritive quality traits: ADF: acid detergent fiber; ADL: acid detergent lignin; DMDig: digestible dry matter; NDF: neutral detergent fiber; NDFD: digestible NDF; Prot: protein; and WSC: water-soluble carbohydrate (right-hand heat ‘columns’). GMR rows and columns are clustered via hierarchical clustering using Euclidean distance as the dissimilarity measure with the ‘complete’ (maximal intercluster dissimilarity) linkage type.

### Statistical models

The dry matter yield records were transformed by applying the natural logarithm scale (base *e*) to correct for the skewed distribution, which is expected in this type of time-series data, and lower the non-constant error variances (heteroscedasticity). Therefore and for simplicity, all models were fitted on the (natural-)log-transformed data [*log(y)*]. For the set of nutritive quality traits, all analyses were performed sustaining the original scale.

In the statistical models described below, phenotypic values of each genotype were regressed on the environmental gradient (EG) or the combination of EG and time points covariables. Thus, random regression coefficients are computed for every entry, hence the model designation. Environments were ordered by mean performance to create the environmental index. Polynomial functions as the basis functions were used to model both longitudinal dimensions. Orthogonal Legendre polynomials (Kirkpatrick et al., 1990) were obtained by first rescaling the time points and environmental means to the range from -1 to 1 using 2.

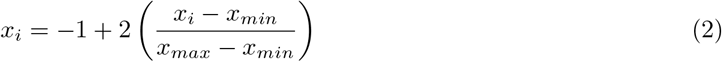

followed by the recursive equation 3.

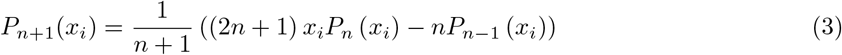

and the normalization according to the equation 4.

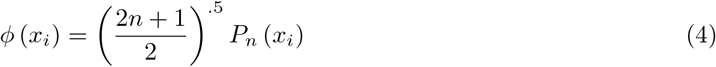

In a matrix setting, Legendre polynomials can be computed as **Φ** = **MΛ**, where **Φ** is a matrix containing the normalized polynomials for harvest time/environment means; **M** store polynomials of standardized harvest time/environment mean values; and **Λ** is the matrix of Legendre polynomial coefficients of order *d* + 1, where *d* is the degree of fit (Schaeffer, 2016). The intercept was set to 1.00 instead of 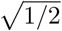 from the full Legendre Polynomials covariates. The choice of the proper polynomial degree for the time and environmental trajectories was made based on assessments of the goodness-of-fit criteria Akaike information criterion - AIC (Akaike, 1974) and Bayesian information criterion - BIC (Schwarz, 1978). The final decision for the appropriate level of flexibility in the model selection process was based on cross-validation, which allows for direct computation of test error as 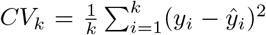. The drop among the estimated test mean squared error for each polynomial fit was computed and the model showing the minimum with the lowest-order degree was selected. Finally, the ultimate random regression models fit Legendre’s polynomial of degree one (linear) over the environmental gradient and of degree two (quadratic) over harvest time. The linear function over the environmental index yields two biologically meaningful coefficients: the intercept (overall performance) and the slope (plasticity) which are random effects that vary freely among individuals.

All models described hereafter contain an additional random term, which is a summation of random effects, that captures the variance due to the spatial field variation as well as border effects. The 2D field layout was scanned in a sliding window manner to capture the spatial variability (see Figure 3). The defined window size covers 14 neighboring experimental units, all equally weighted on the target plot. The sliding window in a given position was unique in each one of the four harvests. A common spatial variance was assumed but the alternative scenario, where environment-specific variances are estimated, is possible to model as shown by Guo et al. (2020).

**Figure 3:**
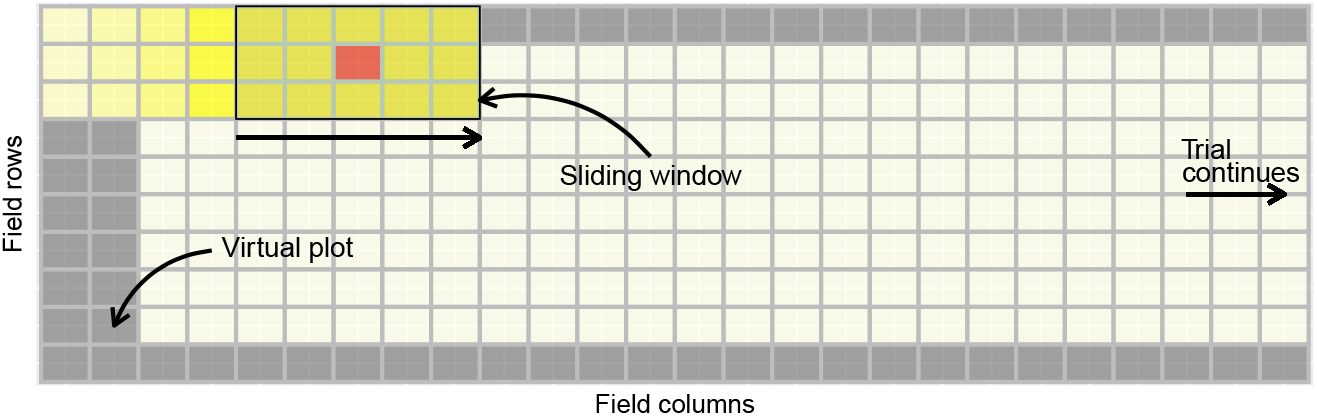
Schematic representation of a segment of one field trial showing the spatial modeling strategy by means of a sliding window over columns within rows to scan for spatial variability. Virtual plots surrounding the trial are highlighted in grey. The moving window takes up 15 plots each time, the targeted plot (red cell) and 14 surrounding plots, to mine for local spatial heterogeneity.

### Random regression baseline model (M1)

The baseline adopted model accounts for the spatial dimension by regressing individuals on environmental gradient Legendre Polynomial covariables. Thus, modeling repeated measurements as an infinite-dimensional (function-valued character) model instead of a character state approach. All four harvest events were summed to compute the total dry matter yield of the whole production year (response vector *y*) and can be seen as a measure of the area under the dry matter production curve. As the nutritive quality traits are expressed as a percentage of DMY, the response vector for each variable was obtained by a weighted average, where the weighting factor was the DMY of each harvest. Therefore, the univariate random regression model in eq. 5 was fitted to the data.

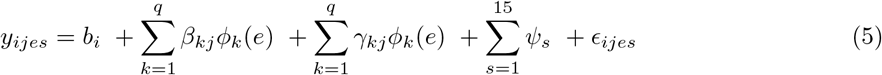

where *y*_*ijes*_ represents the phenotype of each *j* full-sib family at the environment point *e* within the *i*th level of fixed effect *b* (year-location-management classes); *β*_*kj*_ and *γ*_*kj*_ are the *k*th random regression coefficients for the additive genetic and non-additive effects, respectively, for family *j* (the last is referred as permanent environment effect by the animal science community); *ϕ*_*k*_(*e*) represents the *k*th Legendre covariate for the record of family *j* made at the environment point *e*; *q* is the number of covariates; *Ψ* is the spatial effect accounting for 14 neighboring plots plus the target experimental unit (see Figure 3), and *ϵ*_*ijes*_ is the environment-independent random residual for each observation. No fixed regression was fitted once the model term *b*_*i*_ already describes the overall trajectory of the population. The model term capturing the non-additive entry variance over the environmental gradient trajectory can be seen as an overlap of effects, e.g., dominance, epistasis (Kruuk & Hadfield, 2007), and non-random environment effects, that persists throughout the trajectory. As a linear reaction norm model, equation 5 can also be fitted without the use of orthogonal polynomials. In matrix notation, the model specified in 5 can be rewritten as displayed in equation 6.

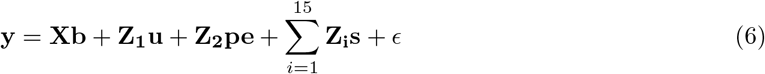

where **y** is the vector of observed phenotypes; **b** is the vector of fixed effects; **u** and **pe** are the vectors of random regressions for additive genetic and non-additive effects, respectively; **s** is a vector of spatial effect; **X** and **Z**_*i*_ are the design matrices linking fixed and random spatial effects to the phenotypic records; **Z**_1_ and **Z**_2_ are covariable matrices containing orthogonal polynomials; *ϵ* is a vector of random residuals. It was assumed a (co)variance structure of the form presented in 7.

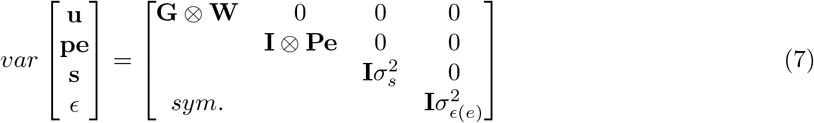

where **G** is the genomic relationship matrix described in equation 1; **W** and **Pe** are 2× 2 (co)variance matrices of random regression coefficients for the additive genetic and non-additive effects, respectively; 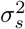 is the spatial effect variance; 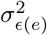 stands for the heterogeneous residual varying for each combination of year-location (four classes in total); **I** is an identity matrix; and ⊗is the Kronecker product. Genomic estimated breeding values (gEBVs) of a family *j* can be computed for each *e* point of the environmental gradient as follows: 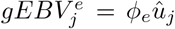, where 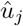 is the 2×1 vector of estimated additive genetic values and *ϕ*_*e*_ is the vector of orthogonal coefficients evaluated at each environment *e*.

### Random regression model (M2)

The baseline model was expanded to allow the modeling for the harvest event. Thus, the linear model in equation 8 accommodates a non-linear function (second-order Legendre polynomial) to model days to harvest and a linear fit over the environmental gradient, allowing the identification of G ×E for combinations of environment and harvest date levels. Therefore, the model is smoothing the covariance matrix by using a reduced fit (*m −* 1).

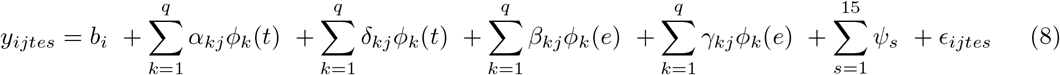

where *y*_*ijtes*_ represents the phenotype of each *j* full-sib family at harvest point *t* and environment point *e* within the *i*th level of fixed effect *b* (year-location-management-harvest classes); *α*_*kj*_ and *β*_*kj*_ are the *k*th random regression coefficients for the additive genetic effects for the *j*th family by harvest and environment classes, respectively; *δ*_*kj*_ and *γ*_*kj*_ are the *k*th random regression coefficients for the non-additive genetic effects for the *j*th family by harvest and environment classes, respectively; *ϕ*_*k*_(*e*) represents the *k*th Legendre covariate for the record of family *j* made at the environment point *e*; *ϕ*_*k*_(*t*) represents the *k*th Legendre covariate for the record of family *j* made at the harvest point *t*; *q* is the number of covariates; *Ψ* is the spatial effect as described in M1; and *ϵ*_*ijtes*_ is the time- and environment-independent random residual for each observation. The matrix representation of the model M2 is as presented in equation 6, with the proper expansion of the order of matrices and vectors to accommodate the harvesting effect. In this model, the combination of year, location, management, and harvest yields 32 subclasses for the fixed part **Xb**. This is further expanded by accounting for the plot forage harvester by date effect (*h/d*), given that each harvest event was generally performed in more than one day and oftentimes with two combines. In addition, the *h/d* of the previous harvest event (except for the first one) was also included. During the data wrangling process, it was noticed a substantial effect of these effects as additional sources of variation. The (co)variance structure of M2 also follows:

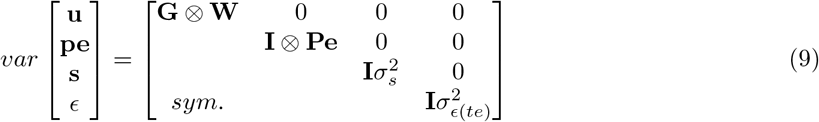

however, the variance-covariance of **u** is now:

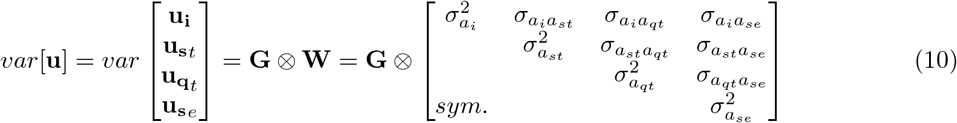

where 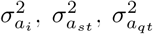, and 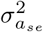 are the additive genetic variances for intercept, harvest slope, harvest quadratic term, and environmental slope, respectively; and **W** is the (co)variance matrix of the additive genetic random regression coefficients. The (co)variance structure for **u** in 10 also applies to *var*[**pe**], however, the genomic relationship matrix **G** is replaced by an identity matrix **I**. Finally, 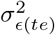 in 9 is the heterogeneous residual variance structure with 16 classes, one for each combination of year-location-harvest.

Genetic parameters were estimated from M2 and include the additive genetic and permanent environmental (co)variance matrices computed as **ΦWΦ**^*′*^ and **ΦPeΦ**^*′*^, respectively. Diagonal elements from these matrix operations are the additive 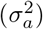 and non-additive genetic 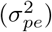 variances for each harvest time or environment class. From the off-diagonal, covariances between *j*th harvest time and *e*th environment were obtained for additive and non-additive genetic effects. The matrix **Φ** of the order *m*·(*n* + 1) contains Legendre polynomials for days to harvest and environmental gradient and takes the following form in M2:

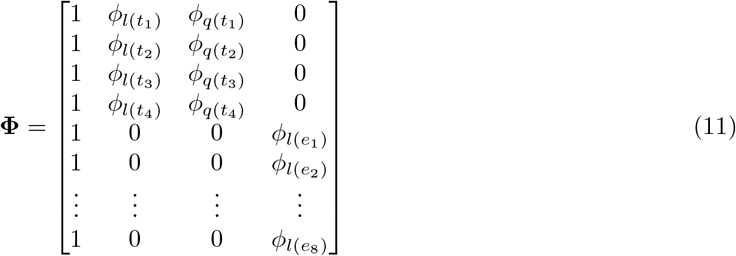

The gEBV of a family *j* from the random regression model can be computed for every combination of time point and environment as 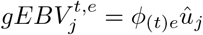 where 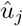 is the 4×1 vector of solutions for family *j*’s additive genetic value and *ϕ*_(*t*)*e*_ is the vector of orthogonal coefficients evaluated in time point *t* and environment *e*. For instance, one can compute the annual DMY for an average environment (predictor equal to zero) by multiplying 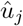 with the first four rows of **Φ** in 11.

Additional genetic parameters were derived from model M2 and it includes estimates of the additive genetic correlation (*ρ*_*a*(*ij*)_) between *i*th harvest time in *j*th level of the environment:

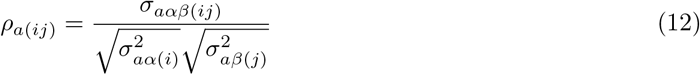

and narrow sense heritability in an entry-mean basis:

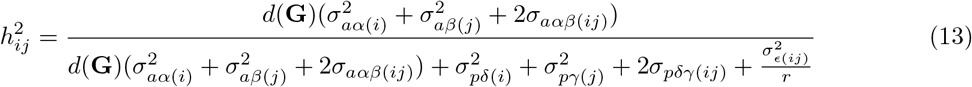

where, *d*(**G**) is the average diagonal element of the G-matrix, 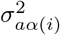 and 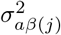 are the trait’s additive genetic variances for the *i* th cut and *j* th environment, respectively; and 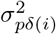 and 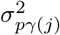 represents the trait’s non-additive variance for the *i* th cut and *j* th environment, respectively; *σ*_*aαβ*(*ij*)_ and *σ*_*pdγ*(*ij*)_ are the additive genetic and non-additive covariances between levels of cut and environment, respectively; 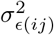 represents the heterogeneous residual variance for each combination of cut and environment; and *r* is the harmonic mean of the number of replicates. Reaction norm trajectories/surfaces were computed for all entries as population-level deviations across time gradient considering an average environment, across the environmental gradient considering an average harvest performance within a production year, and the combination of them using a three-dimensional plot (by forcing the solution vector to be a diagonal matrix). All parameters can be estimated for every combination of harvest and environment within the limit ranges of the study. Genetic parameters were displayed for a total of 16 levels of harvest time considering that the four harvests were not performed exactly the same day in every combination of year-location. Indeed, gRRM can efficiently bookkeep these differences among individuals for levels of covariate indexes.

### Multivariate random regression model (M3)

The multivariate random regression is a more flexible model but at the expense of requiring the estimation of a greater number of parameters. In this setup, the aim was to treat the four repeated harvests as different traits (character-state approach). Again, first-order Legendre polynomials generated using environmental means were regressors in the model for the additive genetic effect and describe the random trajectories of each individual over the environmental gradient. The non-additive random term 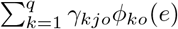 was dropped from the model to reduce the computational burden and to facilitate convergence. Therefore, the model M3 was defined as follows:

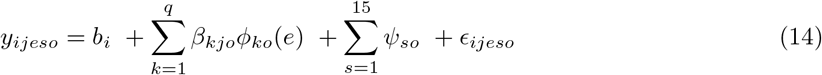

where *y*_*ijeso*_ was the *e*th performance record of the *j*th full-sib family from the *o*th trait (i.e., response variable measured at each harvest); *b*_*i*_ was the fixed effect (year-location-management classes plus *h/d* levels) for the *o*th trait; *β*_*kjo*_ was the *k*th random regression coefficients for the additive genetic effects for family *j*; *ϕ*_*ko*_(*e*) represents the *k*th Legendre covariate for the record of family *j* made at the environment point *e* for the *o*th trait; *q* is the number of covariates; *Ψ* was the spatial effect accounting for 14 neighboring plots plus the target experimental unit (see Figure 3), and assumed independent between traits for simplicity; and *ϵ*_*ijeso*_ is the time- and environment-independent random residual for each observation. In an expanded matrix notation, model M3 can be represented as follows:

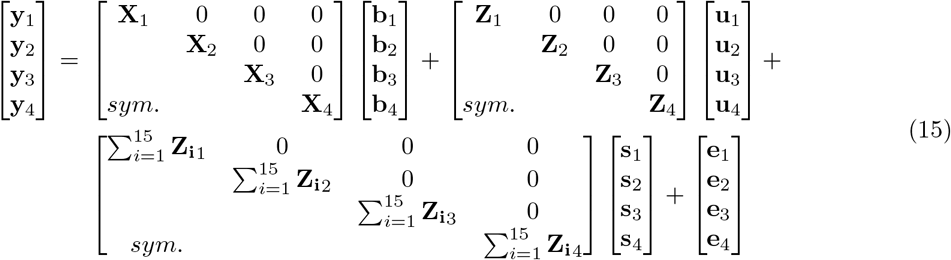

here, **y**_*i*_ are subvectors of phenotype records for trait one to four; **X**_*i*_, **Z**_*i*_, and **Z**_**i***i*_ are incidence matrices linking observations in **y**_*i*_ to the set of fixed effects in **b**_*i*_, random additive genetic effects **u**_*i*_, and random spatial effects **s**_*i*_, at the *i*th measurement time. Variance components were estimated by assuming [**u**_1_ **u**_2_ **u**_3_ **u**_4_]^*T*^ *∼ N* (0, **G** ⊗ **W**), [**s**_1_ **s**_2_ **s**_3_ **s**_4_]^*T*^ *∼ N* (0, **I** ⊗ **S**), and [**e**_1_ **e**_2_ **e**_3_ **e**_4_]^|*T*^ *∼ N* (0, **I** ⊗ **R**). The additive genetic effects for all harvests comes in a 8×8 (co)variance matrix **W** with additive genetic variances for each trait’s intercept and slope on the diagonal and additive genetic covariance between random regression coefficients on off-diagonal positions. It can be shown that if a polynomial of maximum degree (*m −* 1) is used for the temporal dimension on model M2, then estimates of **W** approximates the ones from the full multivariate approach in M3. The covariances between harvests in **S** for spatial and in **R** for residual effects were restricted to zero, while **W** was assumed as unstructured. Heterogeneous residual variances 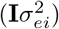 were estimated within harvest for four classes: *i*= 1 to 4 (combination of years and locations). The vector of gEBVs for each *j* family was obtained as 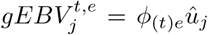, where 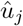 is the vector of solutions for family *j*’s additive genetic value and *ϕ*_(*t*)*e*_ is the vector of orthogonal coefficients for every time point *t* (trait) and environment *e*. The average gEBV was used as e metric of annual performance. Finally, temporal G×E can be assessed from M3 by the magnitude of the additive genetic correlations across time measurements, in which lower values suggests higher interaction.

### Model assessment

The adopted prediction scheme attempts to explore a situation where the goal is to predict the two-year gEBV of newly developed families (unobserved) for an average environment i.e., the specific point of the trajectory where the environmental dimension has zero value. This mimics a scenario where the breeder is interested in assessing the overall adaptability of new breeding material and performing selections based on its rank. Therefore, the validation of genomic-based predictions from the described scheme was performed with a 10-fold stratified cross-validation setup. Genotypes were randomly sampled in a proportional manner within each stratum, represented by the two diallels, and allocated to one of the 10 folds. This ensures that each fold has the same proportion of families from each stratum as the original dataset, therefore, sustaining a similar level of relationship between training and testing sets. Folds constituents were kept constant across prediction models to secure comparability. Hence, for every run, ~90% of the data was used as a training set to predict the remaining masked families in the validation set where phenotypes were discarded. For each model, the ability to predict the genomic breeding value of individuals in the validation (hold-out) set was assessed by the correlation between genomic estimated breeding values of families in the test set from the reduced data (partial dataset) with genomic estimated breeding values from the full dataset (*ρ*_*f,r*_) in which individuals breeding values were predicted using own records (Legarra & Reverter, 2018) and can be seen as a relative measure of prediction accuracy. The quality of fit produced by each model was also assessed by average estimates of bias (intercept *µ*_*f,r*_) and dispersion (slope *b*_*f,r*_) via linear regression of the type *gEBV*_*full*_ = *b*_0_ + *b*_1_ * *gEBV*_*reduced*_ across folds, and the values presented in a table format (Supplemental Table S1). A second study was designed in which the large sparse diallel was used to predict the small one, resembling a scenario where the goal is to use the available information to predict an independent population. Prediction accuracy and quality of fit were assessed as before. Finally, this last study was used to test an additional scenario in which the goal was to forecast the two-year gEBV by recurrently updating model M2 with new measurement data arriving from every new harvest up to the 7^th^, and computing the correlation with the gEBV from the whole dataset in a forward chaining-like cross-validation strategy.

### Computation and Data Visualization

In all described models above, variance components were estimated by average information restricted maximum likelihood (AI-REML) using the DMU package version 6 (Madsen & Jensen, 2013), with convergence precision for REML set to 10^−6^. The DMU4 module was then used for computations regarding all cross-validations. Computations were performed on the GenomeDK high-performance computing facility located at Aarhus University, Denmark. Data wrangling, downstream analyses, and data visualization were performed using the R programing language (R Core Team, 2020). Figures were prepared using ggplot2 (Wickham, 2016), ComplexHeatmap (Gu et al., 2016), and Plotly (Plotly Technologies Inc., 2015) R packages.

## Results

The DMY across cuts follows a typical quadratic curve. The overall DMY of the 1st, 2nd, 3rd, and 4th cuts were 709.9, 360.1, 219.1, and 177.4 g m^2^ for the first production year and 711.1, 153.7, 124.7, and 91.4 g m^2^ for the second, respectively. The fitted model M1 modeled a commonly used response variable conceived by the sum of all cuts within a production year. This model required the estimation of 11 parameters in the maximized likelihood function. When cuts were individually considered, models M2 and M3 were fitted, requiring the estimation of 37 and 56 parameters, respectively, given the imposed restrictions on non-existent covariances on model M3. Despite dropping the random term capturing the non-additive genetic variance in M3 and covariance restrictions, the CPU time necessary to reach convergence was considerably higher compared to the models M2 and M1. Therefore, gRRM M2 is noticeable a more parsimonious model than M3 while fitting simultaneously linear and non-linear reaction norms over a 4x higher dimension of phenotypic data compared to M1. Over the next sections, results of genetic parameters and model comparisons for accuracy of gEBV prediction will be presented and discussed thereafter.

### Spatio-temporal genetic parameters and genomic breeding values

Estimates of variance components, additive genetic correlation, and heritability along the time and environmental gradients were obtained for DMY (Figure 4) and seven nutritive quality traits (Supplemental Figures S1, S2, and S3) by fitting model M2 to the data. The total genetic variance for DMY is mostly explained by the additive term whereas the non-additive (residual genetic) random part of model M2 captures nearly no variance, except in marginal environments. The temporal dimension is the main driver of changes in the DMY additive genetic variance profile while the environmental quality gradient appears to have a constant effect across levels of the time gradient. Similarly, the temporal dimension is the main cause of fluctuation in the additive genetic correlation surface. This conclusion holds also true for all nutritive quality traits (Supplemental Figure S2). In fact, the first cut showed low correlations (≲ 0.50) at all levels of the environmental gradient, which is evidence of strong G×E. It is convened that genetic correlations above 0.8 between environmental exposure variations imply minimum re-ranking of selection candidates whereas estimates below this threshold are evidence of G×E existence (Hayes et al., 2016; Robertson, 1959). Finally, heritability estimates for DMY were generally high across the two-dimensional gradients, with a tendency for increased values in later cuts. However, mostly moderate to low estimates were observed for protein content and WSC (Supplemental Figure S2). The absence of a smooth transition of heritability values across the bi-dimensional gradient space is due to the heterogeneous residual variance accounted for in all fitted genomic random regression models. Overall, heritability estimates suggest sufficient genetic variability to allow breeding gains for all evaluated traits.

**Figure 4:**
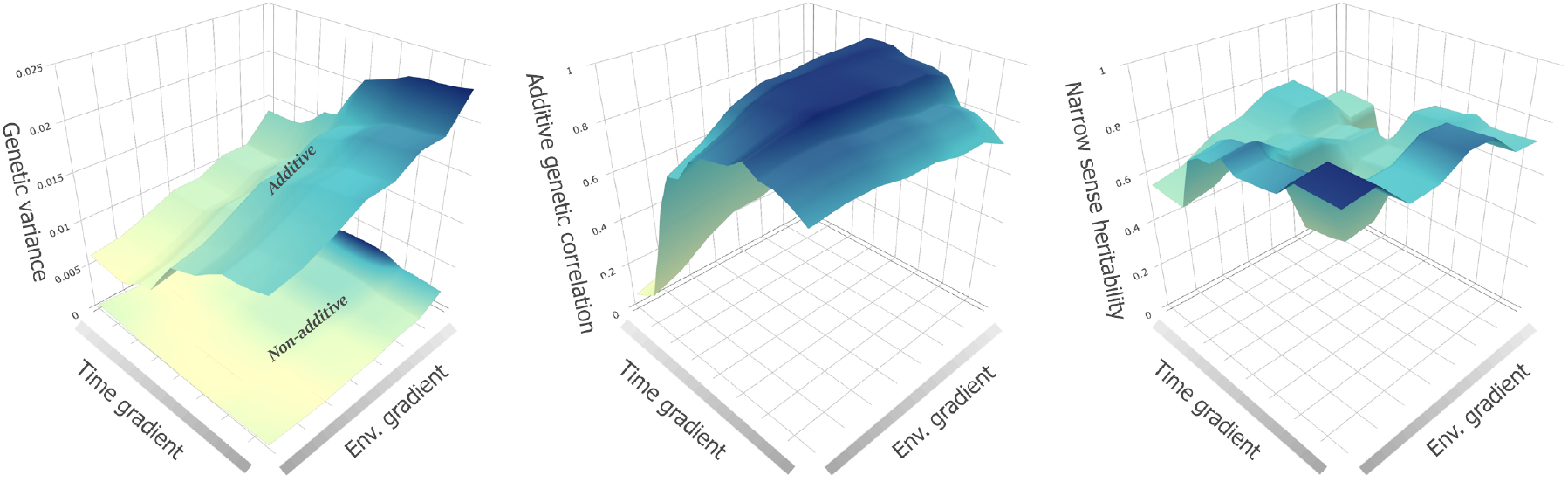
Three-dimensional diagrams showing the functional relationship between estimates of genetic variance (left), additive genetic correlation (middle), and narrow-sense heritability (right), and the two independent variables defined by the temporal and environmental gradients, for the dry matter yield trait.

In addition to the correlations quantified from model M2 and displayed in Figure S4 and Supplemental Figure S2, estimates of genetic correlations derived from multivariate model M3 between pairs of biomass harvests/quality measurements are shown in Figure 5. Overall, there is a tendency for higher correlations between nearby observations, suggesting autocorrelation. This is in accordance with Giri et al. (2019) which reported 0.34 and 0.12 first and second-order temporal autocorrelation, respectively, for ryegrass DMY among harvests. Except for WSC, later cuts show higher correlation values and, consequently, less temporal G×E. The DMY of the first cut (spring harvest) corresponds to nearly half of the total annual biomass mowed/grazed. As Figure 5 shows, there is roughly no association between yield in the first cut and later measurements, implying substantial temporal environment versus genotype interaction. Accordingly, considerably re-ranking of genotypes’ reaction norm curve across levels of time gradient can be observed for the first harvest (Supplemental Figure S4). Besides the bi-dimensional changes in genetic variance and correlation, estimates of the 16 heterogeneous residual variances obtained from model M2 also varied as much as 10 folds (highest-lowest ratio value) along with high correspondence with estimates from model M3 (Pearson = 0.98).

**Figure 5:**
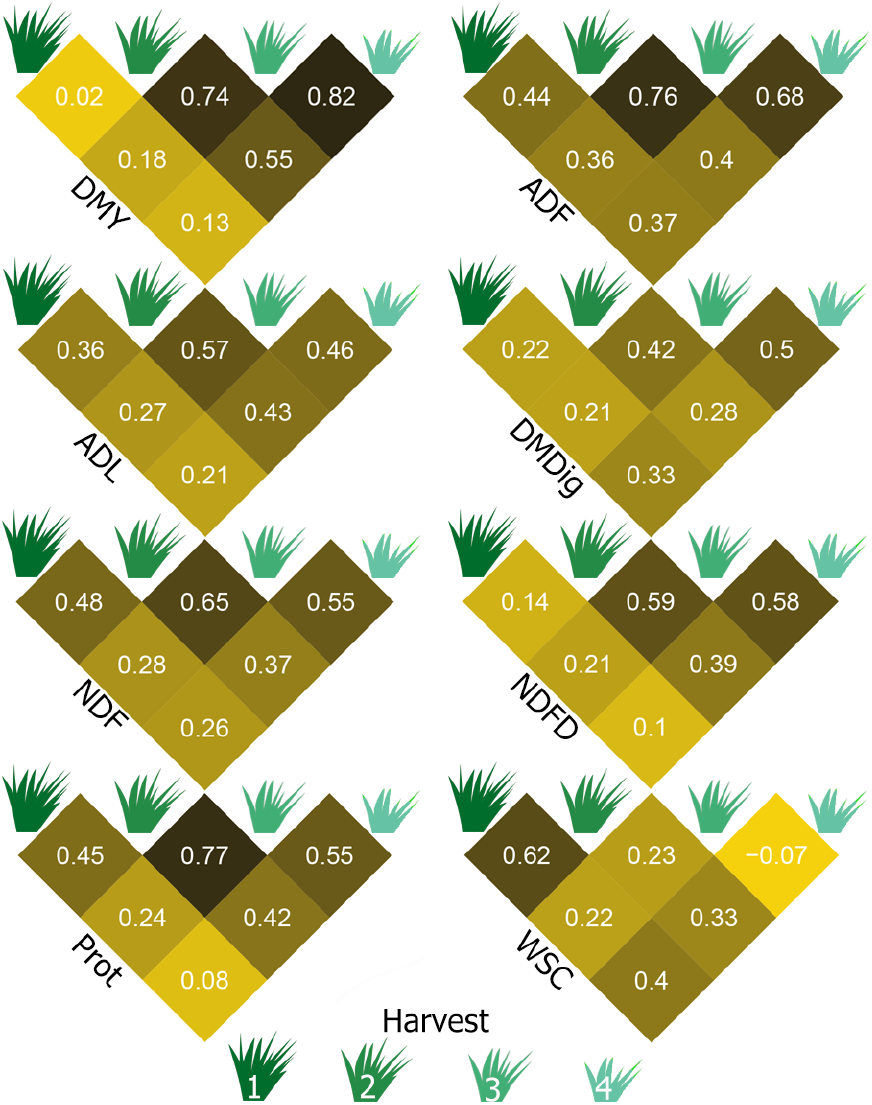
Genetic correlations between pairs of harvests of dry matter yield (DMY) and repeated measurements of nutritive quality traits in perennial ryegrass from an overall environment, estimated via a multivariate random regression model (M3). ADF: acid detergent fiber; ADL: acid detergent lignin; DMDig: digestible dry matter; NDF: neutral detergent fiber; NDFD: digestible NDF; Prot: protein; and WSC: water-soluble carbohydrate.

Fitting the random regression model M2 results in a solution vector of the size equal to four, which can be used to generate surface plots of the reaction norm of every entry, given the appropriate matrices of time and environmental covariables. Figure 6 displays such a surface plot where the top and bottom 20s entries selected by intercept (overall performance) value of DMY are depicted. Complex G ×E in early cuts arises from both time and environmental gradients as surfaces cross each other, entailing re-ranking throughout the two dimensions. In addition, the individual effect of both gradients on the shape of reaction norm trajectories and re-ranking of entries can be seen in Supplemental Figures S4 and S5. Reaction norm slopes from the environmental dimension measure the ability of ryegrass families to respond to increasing environmental quality, assigning them a measure of plasticity which, as the data shows, vary across the temporal spectrum. In fact, slope variance for the second cut from multivariate model M3 was *∼*2.5 larger than of the remaining cuts (data not shown). Also from M3, genetic correlations between intercept and slope varied from -0.24 *±* 0.098 (first cut) to -0.82 *±* 0.032 (second cut), suggesting a certain level of independence for the variance of these two quantities in the first harvest, which leads to an increased possibility of re-ranking over the environmental gradient. Thus, sensibility emerges from both sources of variations as gEBV profiles are not static along gradients.

**Figure 6:**
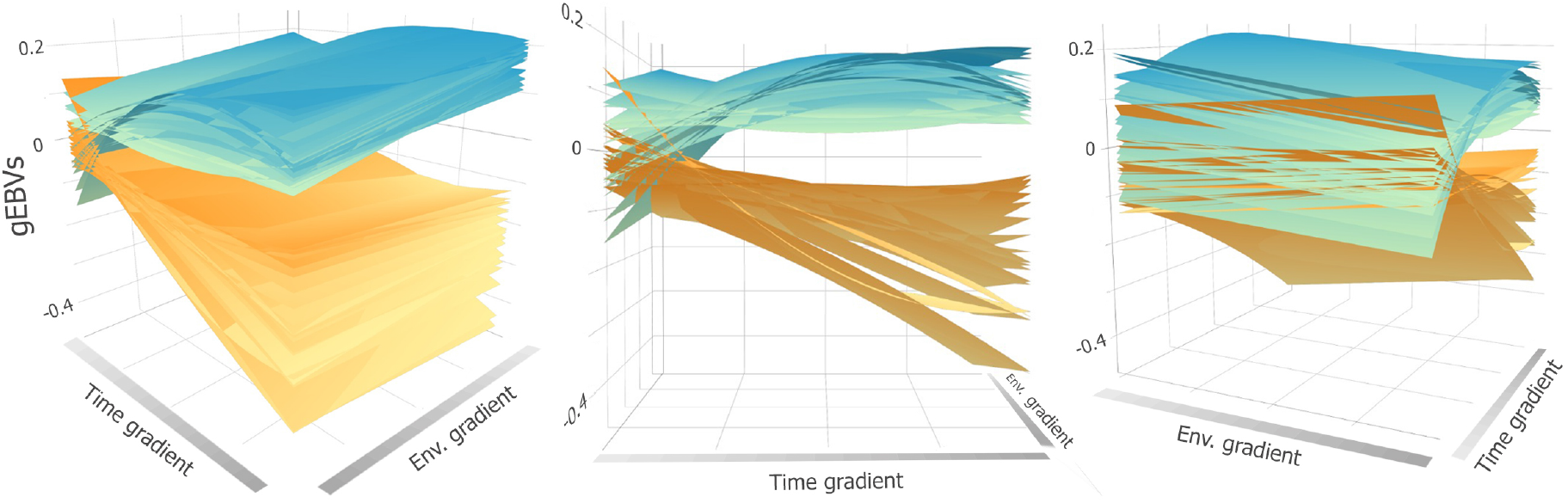
Three views of the 3D surface plot displaying the genomic estimated breeding value (gEBV) profile of the top and bottom 20s ryegrass families by intercept value, estimated via a random regression mixed model for the dry matter yield (DMY) trait. The values on y-axis represent deviations from the population trajectory captured by the fixed effect.

From gRRM model M2, correlation estimates between random intercept and slope were -0.68, -0.46, 0.17, 0.41, -0.73, -0.85, -0.65, -0.11 for DMY, ADF, ADL, DMDig, NDF, NDFD, Prot, and WSC, respectively. These patterns can be seen in Supplemental Figure S5 and are similar to the ones computed using model M1, which only accounts for the environmental dimension (data not shown). As the aforementioned figure shows, little re-ranking across the environmental gradient was observed for traits expressing high correlation between coefficients (NDF and NDFD), intermediate re-ranking for moderate correlation values (DMY, ADF, DMDig, and Prot), and high G×E for the low correlated ones (ADL, WSC). However, no re-ranking was observed between the top and bottom’s 20 families for all traits across the environmental gradient as is displayed in Supplemental Figure S5. If for any of the evaluated traits, genotypes that did better in low-quality environments did worse in high-quality ones and vice versa, were available then high-magnitude negative correlations would be present in Supplemental Figure S2, implying the existence of crossover interaction whithin the range of the environmental covariate. This was not the case given the geographical similarity among environments.

### Genomic Prediction of Breeding Values

#### The whole dataset stratified 10-fold cross-validation

Even though gRRM allows to estimate prediction accuracy throughout the whole trait trajectory, the study focused on the ability to predict entries’ two-year gEBVs by comparing models with and without the recovery of temporal information. Predicting genomic estimated breeding values in related populations yielded, as expected, high magnitude prediction accuracy values (Figure 7). The average Pearson correlation coefficients across traits and models were considered high (*>*0.75), except for WSC (*∼* 0.60) using model M1. Prediction accuracy was slightly larger for model M1 when predicting dry matter yield and digestibility. An increase in accuracy for WSC and in lower magnitude for protein and the fiber variables (NDF and ADF) was due to the inclusion of the temporal dimension simultaneously with environmental classes into the statistical models M2 and M3. The breeding values of nutritive quality traits ADF, NDF, and WSC (M2 and M3) were easier to predict than the others. The values of the regression coefficients (*µ*_*f,r*_ and *b*_*f,r*_) which reveal bias and inflation or deflation (over or under-dispersion) are available in Supplemental Table S1. For bias, whose values *µ*_*f,r*_ *<* 0 underestimate and *µ*_*f,r*_ *>* 0 overestimate predicted gEBVs, results shows only slight deviations from zero, indicating unbiasedness of all model-trait combinations. On the other hand, values of slope (*b*_*f,r*_) were slightly superior to one, indicating a low level of deflation for all three models. At this point, it is relevant to mention that these semi-parametric estimates of fitting quality were improved by considering heterogeneous residual variances.

**Figure 7:**
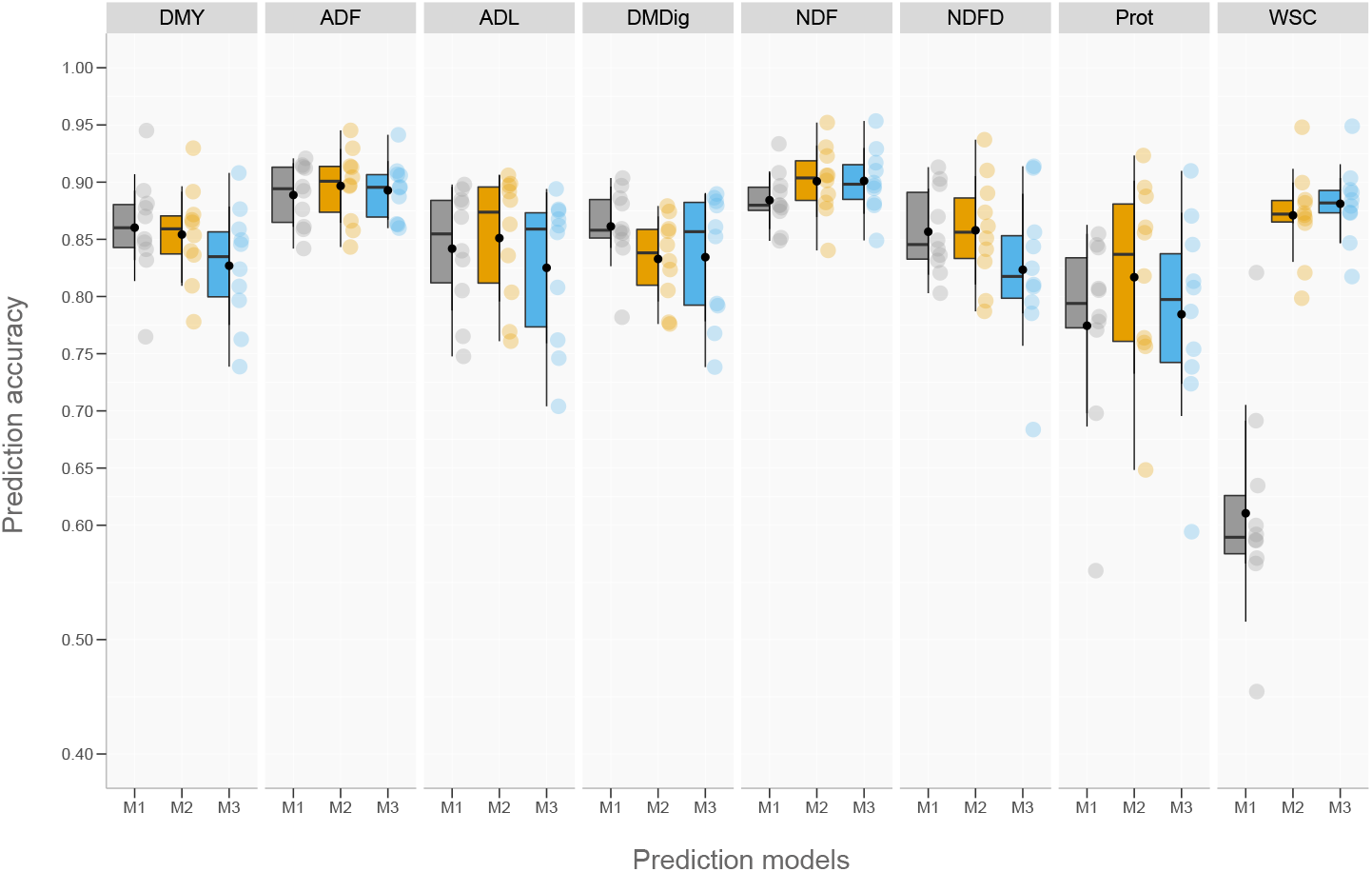
Correlation-based prediction accuracies of genomic estimated breeding values calculated by fitting three predictive models in a stratified 10-fold cross-validation procedure to phenotypic data of eight ryegrass traits. DMY: dry matter yield; ADF: acid detergent fiber; ADL: acid detergent lignin; DMDig: digestible dry matter; NDF: neutral detergent fiber; NDFD: digestible NDF; Prot: protein; and WSC: water-soluble carbohydrate.

#### Genomic prediction in an independent population

No known information from relatives is shared between training and testing sets for the cross-validation scheme depicted in Figure 8. The genetic relationships were captured by dense markers in identity-by-state (IBS) between subsets of individuals. In this scenario, a substantial drop in prediction accuracy was observed compared to the whole dataset cross-validation scheme in Figure 7, especially for ADL, DMDig, and NDFD. A clear separation of the prediction accuracy among the three tested models for DMY shows model M1 with a 12% higher correlation compared with multivariate model M3. Despite the decreased accuracy, model ranking did not change for the majority of assessed traits when comparing scenarios depicted in Figure 7 and Figure 8. The absence of a close relationship between training and test sets when using diallel A to predict diallel B drove poor metrics of bias and dispersion when predicting gEBVs. Estimates of bias from models M2 and M3 somewhat diverged from zero when evaluating nutritive quality traits, except for ADL and Prot, suggesting a tendency for wrong estimates of genetic trends. In addition, slope values deviated from one, indicating the existence of over and under-dispersion of the estimates. In general, the inclusion of the temporal dimension decreases slope values in favor of over-dispersion.

**Figure 8:**
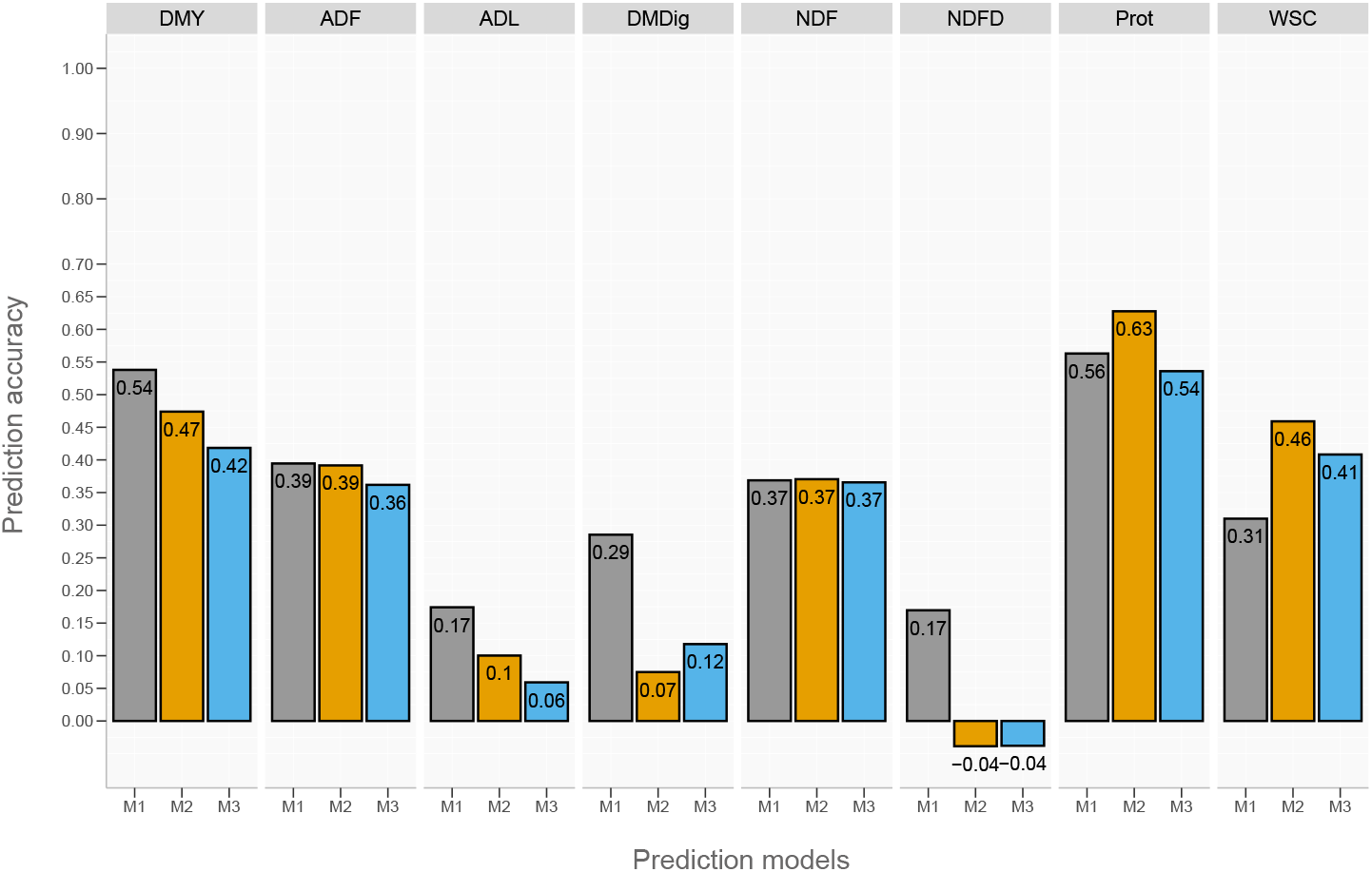
Correlation-based prediction accuracies of genomic estimated breeding values calculated by fitting three predictive models to phenotypic data of eight ryegrass traits. The training and testing sets were defined by the larger and smaller diallel populations, respectively. DMY: dry matter yieldADF: acid detergent fiber; ADL: acid detergent lignin; DMDig: digestible dry matter; NDF: neutral detergent fiber; NDFD: digestible NDF; Prot: protein; and WSC: water-soluble carbohydrate.

#### Longitudinal genomic prediction

Here, it is explored the usefulness of updating the predictive model M2 with data coming from new harvests in a forecasting task. The compelling part of Figure 9 lies on the left-hand side of the plot before the accuracy plateau is reached. Moving from position zero, whose values are the same as in Figure 8, to one on the x-axis implies feeding the model with the first harvest measurements, which differently affects the prediction power of two-year gEBVs. In this scenario, a moderate improvement was observed for DMY and WSC, no gain in accuracy was observed for protein, whereas high improvements were detected for the remaining traits. Adding the second harvest data substantially improves the model capability in predicting the two-year DMY and WSC gEBVs. The magnitude of prediction accuracy estimates scaled fast from low and moderate values in the situation where no identity-by-descent (IBD) between training and testing sets as well as no phenotypic information of entries in the test set (zero on the x-axis) were present, to further high values when predictions were conditioned to the opposite scenario. The results offer evidence for the ability of random regression models to accurately predict breeding values in advance by leveraging early cutting data.

**Figure 9:**
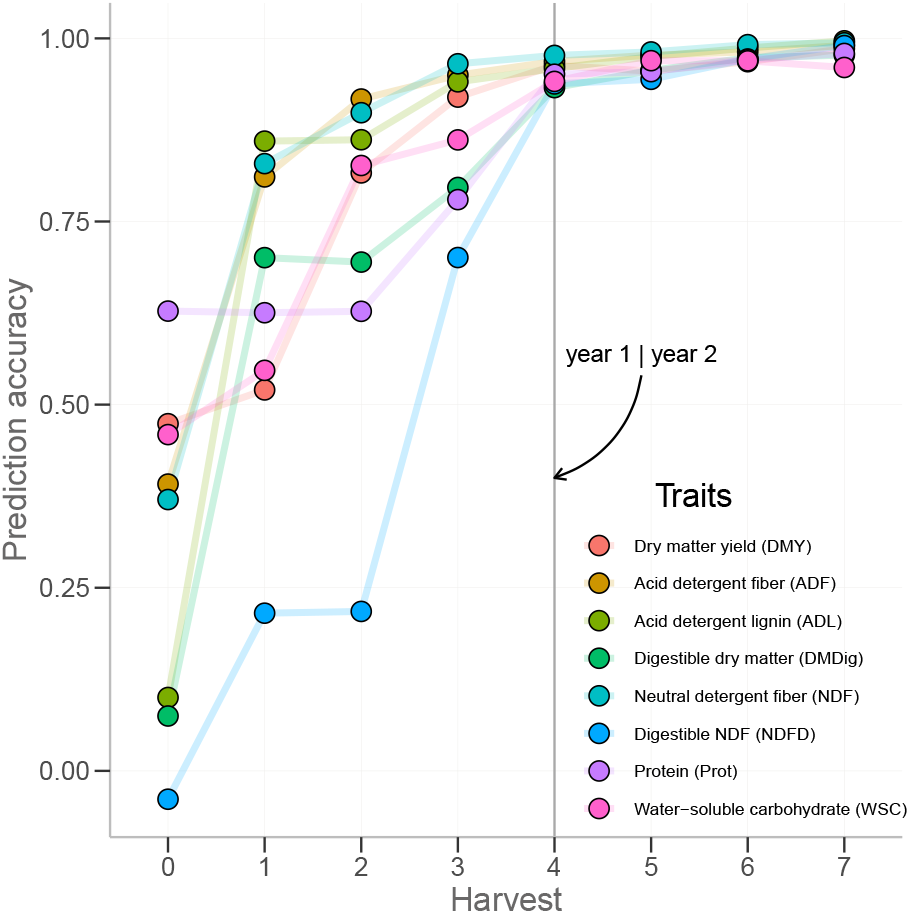
Prediction accuracies of genomic estimated breeding values calculated in a forward chaining-like cross-validation strategy. The training and testing sets were defined by the larger and smaller diallel populations, respectively. At each round of the cross-validation, the random regression model was updated with new harvest data up to the seventh.

## Discussion

In this study, reaction norms were constructed by regressing phenotypes on orthogonal polynomial covariates defined over an environmental quality gradient (single dimension, linear fit) and by simultaneously considering the environmental and temporal gradients (two dimensions) in a multi-environment, multi-harvest setting via alternative random regression modeling approaches. Despite its attractiveness, plant biologists have seldom used random coefficient models for measuring variation in reaction norms in studies of phenotypic plasticity (Arnold et al., 2019). It has an appealing application for the study of complex and economically important traits of forage grasses given the longitudinal nature and the fact that repeated measurements do not totally reproduce the same trait. Repeated measurements from the same genotype over time are subjected not only to a stronger change in the environmental conditions but also to age-related changes and the impact of sequential harvests of aboveground biomass. This may cause more changes to the landscape of gene expression and consequently phenotyping expression than measurements recorded at the same plant growth stage across locations. This complex (co)variance structure can be well modeled using gRRM as shown in the present study. Likewise, the spatio-temporal observed changes in trait variance imply the necessity for heterogeneous residual variance modeling. Accounting for heterogeneous residual variances in gRRM yields better goodness-of-fit estimates (Moreira et al., 2021; Brito et al., 2017). Additionally to other sources of local environmental variations within a field trial, spectroscopy-based traits, as well as DMY, are dependent on the water content of the harvested material, which varies substantially within a cutting day. This spatial variation may not be fully accounted for by prior blocking strategies and downstream corrections, leading to significant spatial inter-dependencies. Hence, all fitted models accounted for spatial heterogeneity by an equally weighted moving average grid which had a light effect on computer memory and CPU load. The possibility of accounting for all the foregoing sources of variation in a classic mixed model framework as well as allowing curvatures in the shape of the reaction norm motivated the use of gRRM models to the analysis of ryegrass multi-environment test data.

The mean performance obtained in a given environment can be seen as the product of all interacting factors conditioning genetics to a certain overall expression landscape. Therefore, it is the ultimate single measure to describe that environment and can be used to construct an index to explain phenotypic plasticity exhibited by crops. In this study, the spatial dimension did not arise from typical plant breeding multi-environment trials (MET) but instead, the limited number of locations was expanded by accounting for nitrogen management and years of evaluation. Therefore, metrics characterizing G×E interaction could be estimated at trait-level and include slope and intercept-slope correlation. The first is a measure of sensitivity to changes in the environment whereas the latter defines the degree of association between performance and sensitivity. The -0.68 intercept-slope correlation estimated for DMY implies that as ryegrass families are exposed to superior environmental conditions the genetic variance becomes smaller. This was also the case for fiber traits and protein. This may be an artifact of the defined environmental gradient, which has a somewhat limited number of evaluation test sites and high similarity among them. It may also be due to changes in weather conditions across years. Overall, moderate G×E interaction was detected for DMY which is in line with other studies evaluating ryegrass across multiple sites (Fè et al., 2015; Conaghan et al., 2008).

As annual plant growth and development progressed and measurements were performed, a temporal gradient was defined. As measurement events slid over this index, the DMY additive genetic variance increased (Figure 4). This can be attributed to the within-year differential mid-season to autumn plant growth among breeding materials, increasing genotypic variance in later cuts of perennial ryegrass as also reported in other studies (Arojju et al., 2020; Fè et al., 2015). In fact, later growth periods were found as the ones with the highest breeding gain for ryegrass DMY over the last decades whereas no improvement was detected for the spring cut (Sampoux et al., 2011). Similarly, variation in the extent of entries’ persistence between years may explain the reason why additive genetic variance was higher in lower DMY environments as it was defined by second-year’s average performances across locations and managements. However, the literature is not clear on the actual existence of a significant genetic component on perennial ryegrass persistence (Dodd et al., 2018; Faville et al., 2020) and no genetic gain was detected for this trait during the last decades (McDonagh et al., 2016). Large genetic correlations (*>*0.80) between DMY with origin on the second to the fourth cuts and DMY from all levels of the environmental gradient implies the possibility of selecting for high genetic value in middle and later cuts aiming to exploit the correlated response across environments. Therefore, selection for DMY and nutritive quality traits can be benefited by assigning a limited weight to the spring cut performance.

The effect of spatio-temporal gradients on genotypic plasticity was demonstrated to be largely additive for all assessed traits. Guo et al. (2018) reported a high proportion of additive to the total genetic variance for dry matter yield of tetraploid ryegrass when using a sufficient larger set of SNPs (80 to 100k). The near-zero residual genetic variance found in the present study can be due to an improved modeling of G and G×E as random intercepts and slopes were fitted, reducing confounding between the two terms, which can occur when only random intercepts are fitted. Considering all traits, the spring cut was poorly genetically correlated with all later cuts and across all levels of the environmental gradient, especially in high yield environments (first production year across locations and managements). The same correlation pattern, along with higher heritability in later cuts, was reported for DMY by Fè et al. (2015) evaluating data from a commercial breeding program of perennial ryegrass. Besides higher genetic variance and genomic heritability in later cuts, Arojju et al. (2020) also reported higher values of predictive ability compared to the estimates obtained for spring harvests.

Variations in environmental features driving genetic responses and shaping phenotypic expression across a continuous gradient also condition gEBVs to re-ranked states. The intensity of G×E was trait dependent, with moderate importance for dry matter yield over the spatial dimension range defined in this study. On the other hand, it was detected a high to low likelihood of re-ranking from the spring to the autumn cuts. Mixed results were found for the remaining traits (Supplemental Figure S4). Given the latitude of the test sites (above the 53rd parallel north), total solar irradiance and average temperature approximate a bell-shaped curve, peaking during the summer. These yearly temporal changes on climate parameters affect not only dry matter yield over successive cuts but also nutritive quality parameters (Loaiza et al., 2016), explaining the significant variation of the quadratic term and the entries’ curvature of reaction norms over the temporal gradient. In fact, a polynomial fit of the seasonal growth of perennial pasture appears to be a reasonable approach to describe longitudinal variations across that continuum (Demanet et al., 2015; Jensen et al., 2014).

Increasing model flexibility aiming at higher accuracy usually leads to less interpretable models, a condition known as the accuracy-interpretability trade-off. Besides decreasing model interpretability, higher-order polynomials can overfit the training data, resulting in high prediction variance. In the present study, genotypes’ response to changes in the environmental gradient was chosen to be a linear reaction-norm, allowing easy interpretation of the slope-coefficient as a plasticity estimate. The temporal dimension showed a curvature where a quadratic term was needed to capture the true relationship more accurately according to the model selection criteria. In biological terms, it can be argued that gRRM models can depict the genetic mechanism of quantitative traits which are products of genes being switched on and off across time and environmental gradients (Yin et al., 2014). Results of these models can be presented in different ways e.g., one point on the curve or one of the curve parameters (Schaeffer, 2016). Here, it was presented the total gEBV of a production year for an overall environment to allow comparisons among the three models. Overall, results offer evidence that modeling non-linearities of phenotype expression over time leads to the improved predictive ability of reaction norm linear mixed models for ryegrass nutritive quality traits. The results presented in this study showed gRRM M2 as a superior modeling strategy to the multitrait approach both in terms of computational requirements and ability to predict gEBVs. On the other hand, the accuracy of gEBV prediction for DMY was favored by the simplest reaction norm model (M1), especially when training and testing sets were unrelated. Therefore, the first-order reaction norm model seems to be advantageous if the goal is to predict the total DMY of a production year. If the seasonal genetic evaluation is needed, then the flexibility offered by gRRM model M2 would be preferred. Besides advantages as environmental correction on a harvest-date basis and flexibility regarding phenotype recording, the random regression model is also suitable for early evaluation of genetic merit in prediction settings as also pointed out by Campbell et al. (2018), which can be useful for reducing the breeding cycle. The forecasting task of future breeding values when plants are already in the field is of great utility as it can reduce the recurrent selection interval and, as a consequence, accelerate the rate at which favorable alleles are combined in new offspring. In addition, it allows breeders to infer about the future instead of waiting until the end of the crop year to look backward.

The breeding goal of improving ryegrass nutritive quality is more relevant than ever and is crucial for sustainable animal production in the face of climate change. Besides dry matter yield per unit of area, improving dry matter digestibility, among other quality parameters, can lead to increased forage intake and use (Byrne et al., 2018). By leveraging random coefficient models, breeders can better investigate the seasonal and spatial changes in genetic variability for target traits and better infer about differences among selection candidates in multiple harvest-location crops. A rearrangement of factors analyzed in this study may be advantageous for a practical breeding aspect, where many more locations of tests are usually available. In the multivariate random regression setting, years can be considered as correlated traits at the same time as trait trajectories are modeled over harvest time and environmental gradients. Further improvements may also be achieved by the inclusion of additional environmental explanatory features such as weather and soil data, allowing deeper investigations into the multidimensionality of phenotypic responses and to perform predictions of individuals in unobserved environments. Finally, the powerfulness of gRRM is also of relevance in the context of plant phenomics, which produces large-scale high-resolution spatio-temporal data, which requires proper statistical models and often the integration of next-generation sequencing data.

## Conclusions

Field evaluation of multi-harvest plant species in multi-environment is resource-intensive, thus appropriate statistical methodologies are required to make sense of all generated data. In this study, it is shown that gRRM models are a robust statistical approach to deal with data coming from perennial ryegrass research, yielding biological meaningful coefficients and high accuracy of gEBV prediction, especially in related training-testing prediction settings. Increasing model complexity by including harvest-level data into reaction norm models appears to improve the ability to predict breeding values for herbage nutritive quality parameters but not for two-year breeding value prediction of dry matter yield.

## Abbreviations

ADF: acid detergent fiber
ADL: acid deterged lignin
DMDig: dry matter digestibility
DMY: dry matter yield
DNBseq: DNA nanoball sequence technology
EG: environment gradient
G×E: genotype-by-environment interaction
GBS: genotype-by-sequence
gEBV: genomic estimated breeding value
GMR: genomic relationship matrix
gRRM: genomic random regression model
NDF: neutral deterged fiber
NDFD: digestible NDF
Ne: effective population size
PCA: principal component analysis
Prot: protein
REML: restricted maximum likelihood
SNP: single nucleotide polymorphism
WSC: water-soluble carbohydrate.

## Acknowledgments

This project was carried out as part of the GreenSelect project funded by GUDP, grant no 34009-15-0952, and the Breed4Biomass project (Innovation Fund Denmark, grant 6150-00020B). The authors gratefully acknowledge the financial support provided by the aforementioned organizations.

## Author contributions

EB designed and led the study, performed the statistical analyses, and wrote the original draft, and included the revision from all authors. DF contributed with plant material development, field trial design, data wrangling, and interpretation of the results. IL contributed with bioinformatics pipelines. MG contributed to the planning, execution, and evaluation of field trials.TD contributed to the calibration and assessment of NIR-based nutritive quality traits. CSJ contributed to the conception of the GreenSelect project and plant material development. TA contributed to the conception of the GreenSelect project and the interpretation of the results. LJ contributed to the conception of the GreenSelect project, provided statistical support, and contributed to the manuscript writing and revision. All authors read and approved the final manuscript.

## Supplemental Material

### Supplemental figures

**Figure S1:**
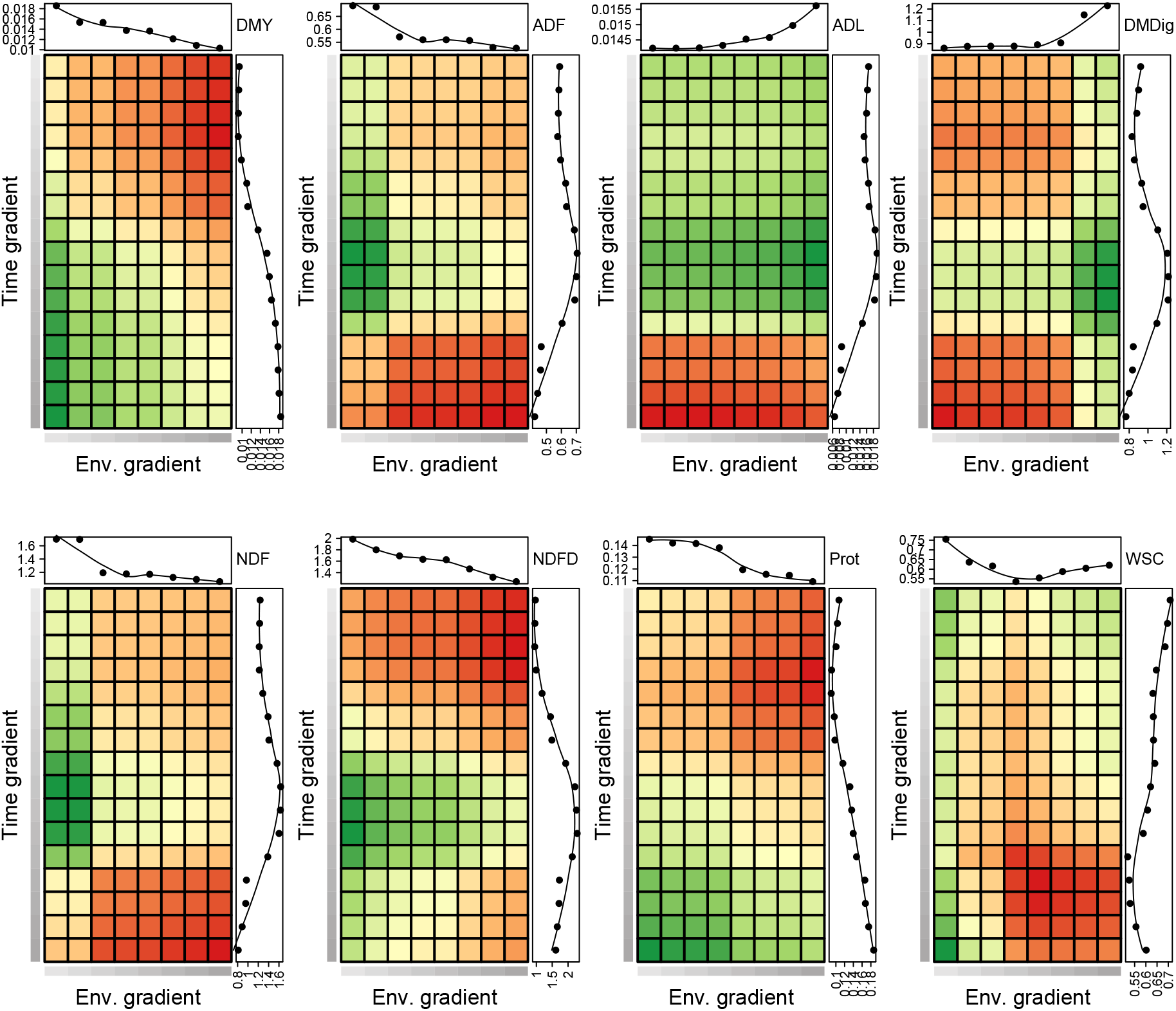
Additive genetic variance 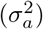 between each combination of time and environmental points, estimated with a random regression mixed model for eight ryegrass traits. Each subplot depicts higher values of 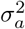 in green whereas lower values are portrayed using red. The intensity of grey color on bars to the left and bottom positions of each subplot depicts the gradients from low values (brighter color) to higher values (dark grey). The line plot graphics on the top and right-hand sides are annotations showing the average 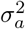 of all time points in a given environment and all environmental points in a given level of harvest time, respectively. DMY: dry matter yield; ADF: acid detergent fiber; ADL: acid detergent lignin; DMDig: digestible dry matter; NDF: neutral detergent fiber; NDFD: digestible NDF; Prot: protein; and WSC: water-soluble carbohydrate.

**Figure S2:**
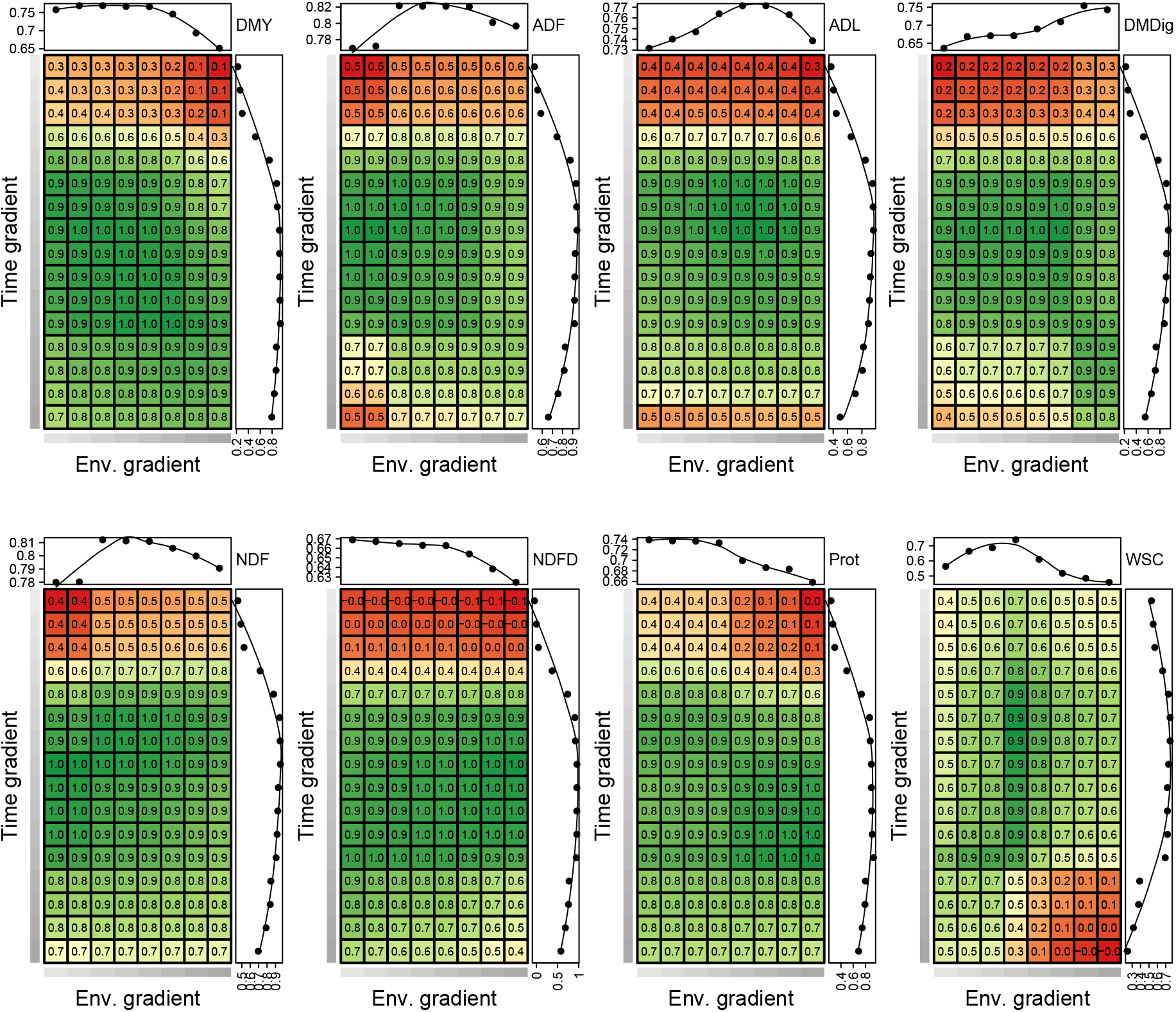
Additive genetic correlation (*ρ*_*a*_) between each combination of time and environmental points, estimated with a random regression mixed model for eight ryegrass traits. Each subplot depicts higher values of *ρ*_*a*_ in green whereas lower values are portrayed using red. The intensity of grey color on bars to the left and bottom positions of each subplot depicts the gradients from low values (brighter color) to higher values (dark grey). The line plot graphics on the top and right-hand sides are annotations showing the average *ρ*_*a*_ of all time points in a given environment and all environmental points in a given level of harvest time, respectively. DMY: dry matter yield; ADF: acid detergent fiber; ADL: acid detergent lignin; DMDig: digestible dry matter; NDF: neutral detergent fiber; NDFD: digestible NDF; Prot: protein; and WSC: water-soluble carbohydrate.

**Figure S3:**
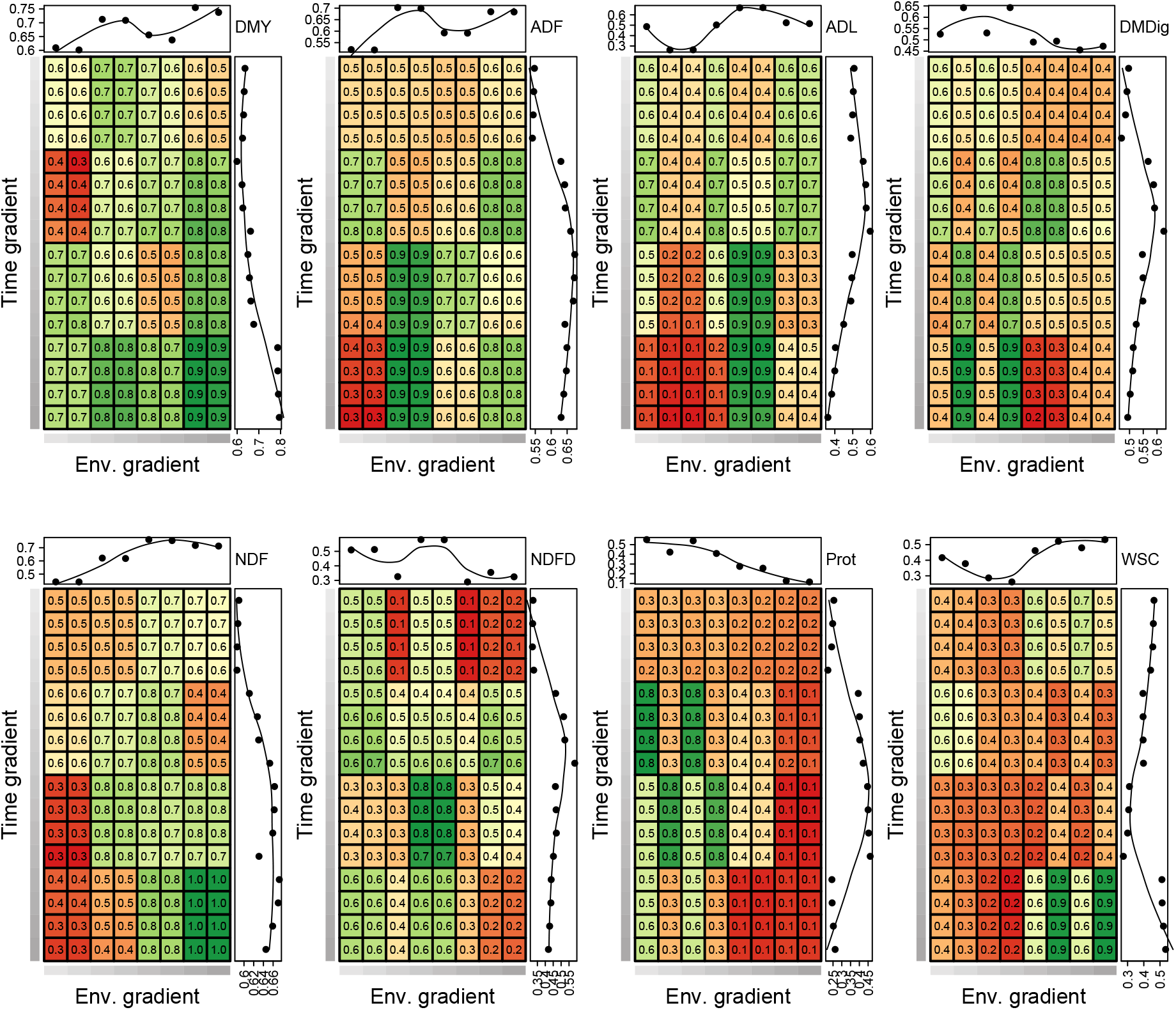
Narrow-sense heritability in an entry-mean basis (*h*^2^) between each combination of time and environmental points, estimated with a random regression mixed model for eight ryegrass traits. Each subplot depicts higher values of *h*^2^ in green whereas lower values are portrayed using red. The intensity of grey color on bars to the left and bottom positions of each subplot depicts the gradients from low values (brighter color) to higher values (dark grey). The line plot graphics on the top and right-hand sides are annotations showing the average *h*^2^ of all time points in a given environment and all environmental points in a given level of harvest time, respectively. The absence of a smooth transition along the gradients is due to the heterogeneous residual variances. DMY: dry matter yield; ADF: acid detergent fiber; ADL: acid detergent lignin; DMDig: digestible dry matter; NDF: neutral detergent fiber; NDFD: digestible NDF; Prot: protein; and WSC: water-soluble carbohydrate.

**Figure S4:**
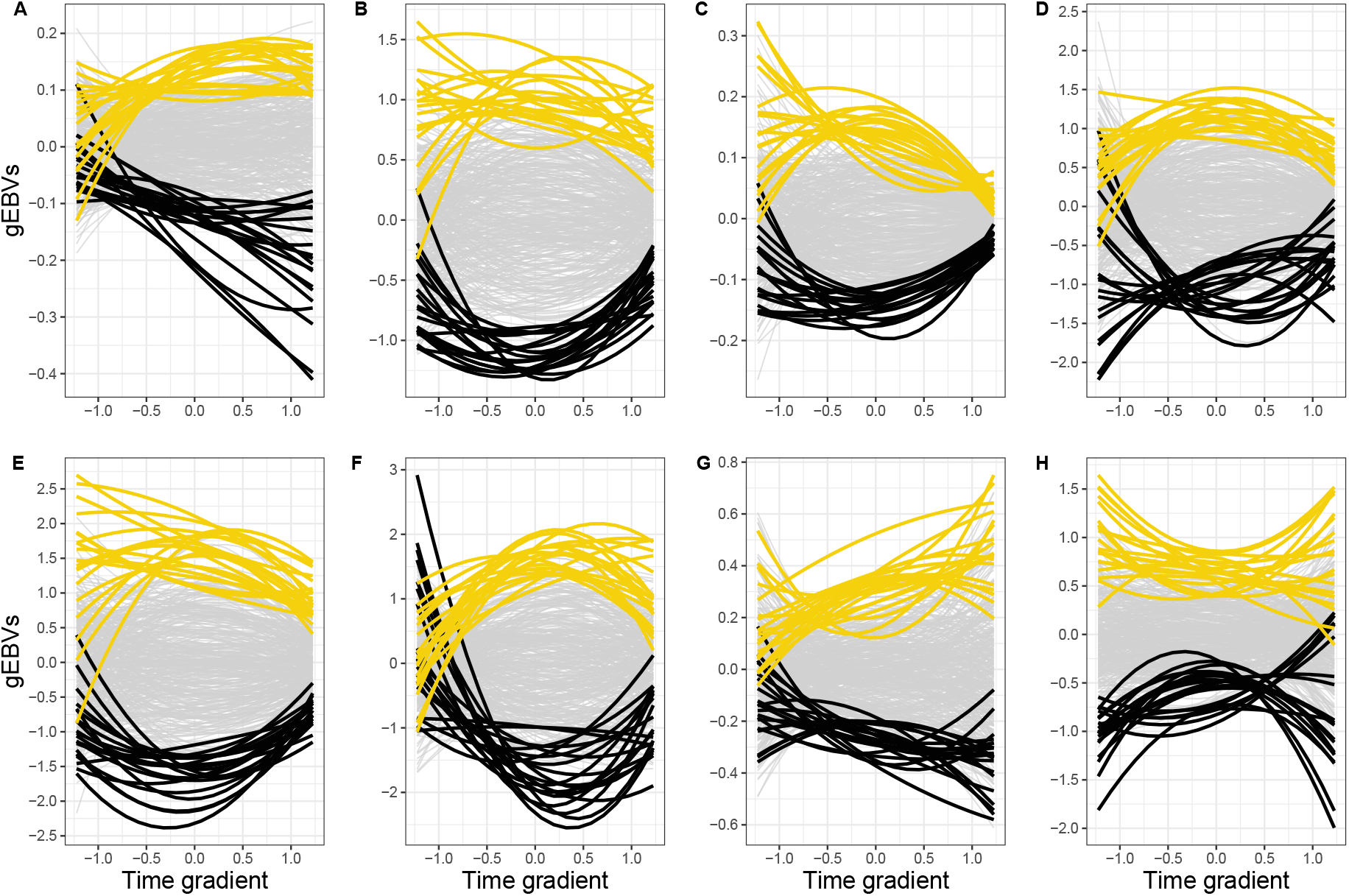
Families’ plasticity is depicted as a reaction norm on different levels of time to measure a given trait along a temporal gradient for on overall environmental. Norms of reaction from genomic estimated breeding values (gEBVs) are highlighted for the top (yellow) and bottom (black) 20 F_2_ families by the intercept value. A: dry matter yield; B: acid detergent fiber; C: acid detergent lignin; D: digestible dry matter; E: neutral detergent fiber (NDF); F: digestible NDF; G: protein; H: water-soluble carbohydrate. The values on y-axis represent deviations from the population trajectory captured by the fixed effect.

**Figure S5:**
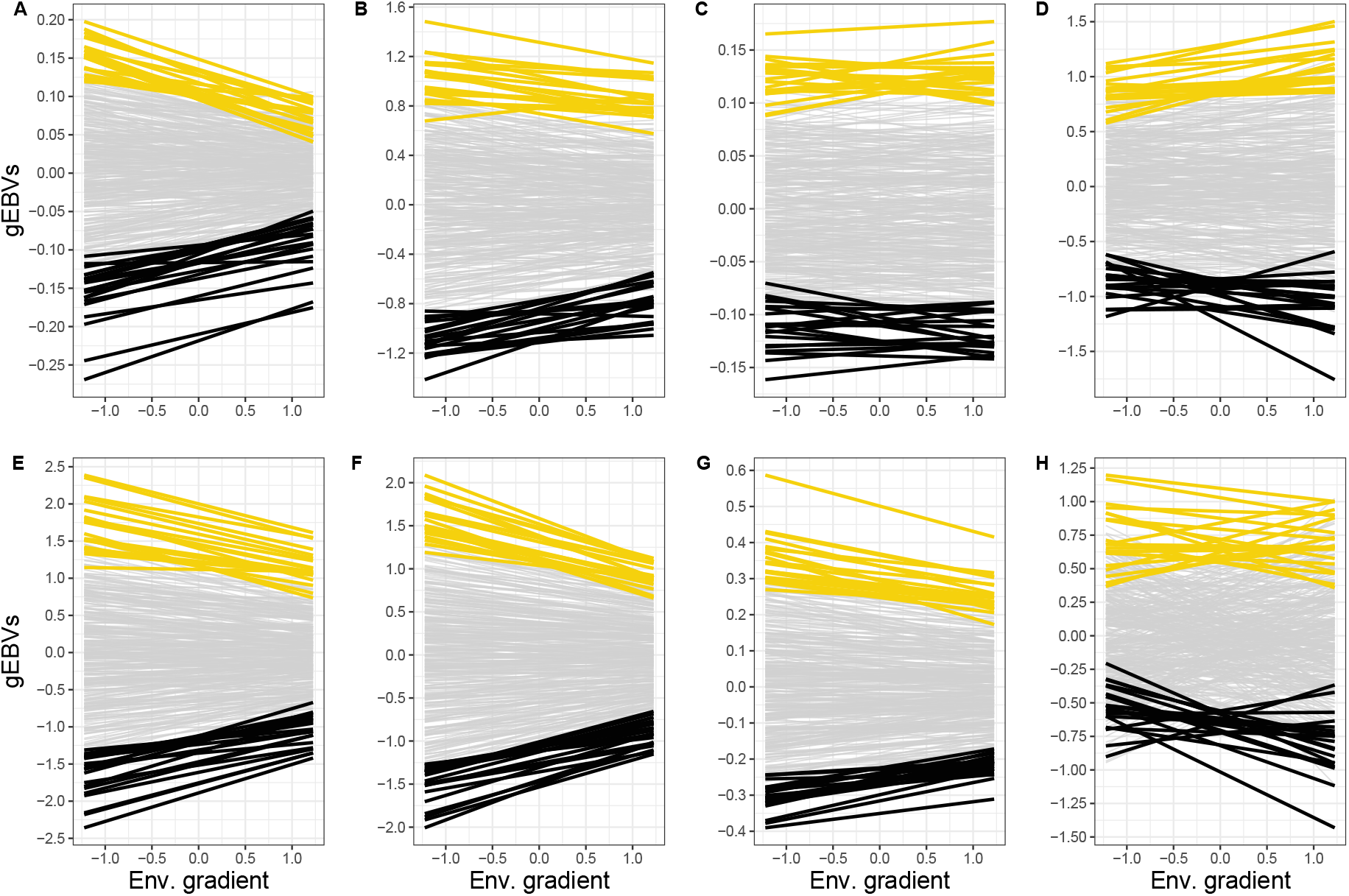
Families’ plasticity is depicted as a reaction norm on different levels of an environmental gradient for an overall harvest performance. Norms of reaction from genomic estimated breeding values (gEBVs) are highlighted for the top (yellow) and bottom (black) 20 F_2_ families by the intercept value. A: dry matter yield; B: acid detergent fiber; C: acid detergent lignin; D: digestible dry matter; E: neutral detergent fiber (NDF); F: digestible NDF; G: protein; H: water-soluble carbohydrate. The values on y-axis represent deviations from the population trajectory captured by the fixed effect.

### Supplemental tables

**Table S1:**
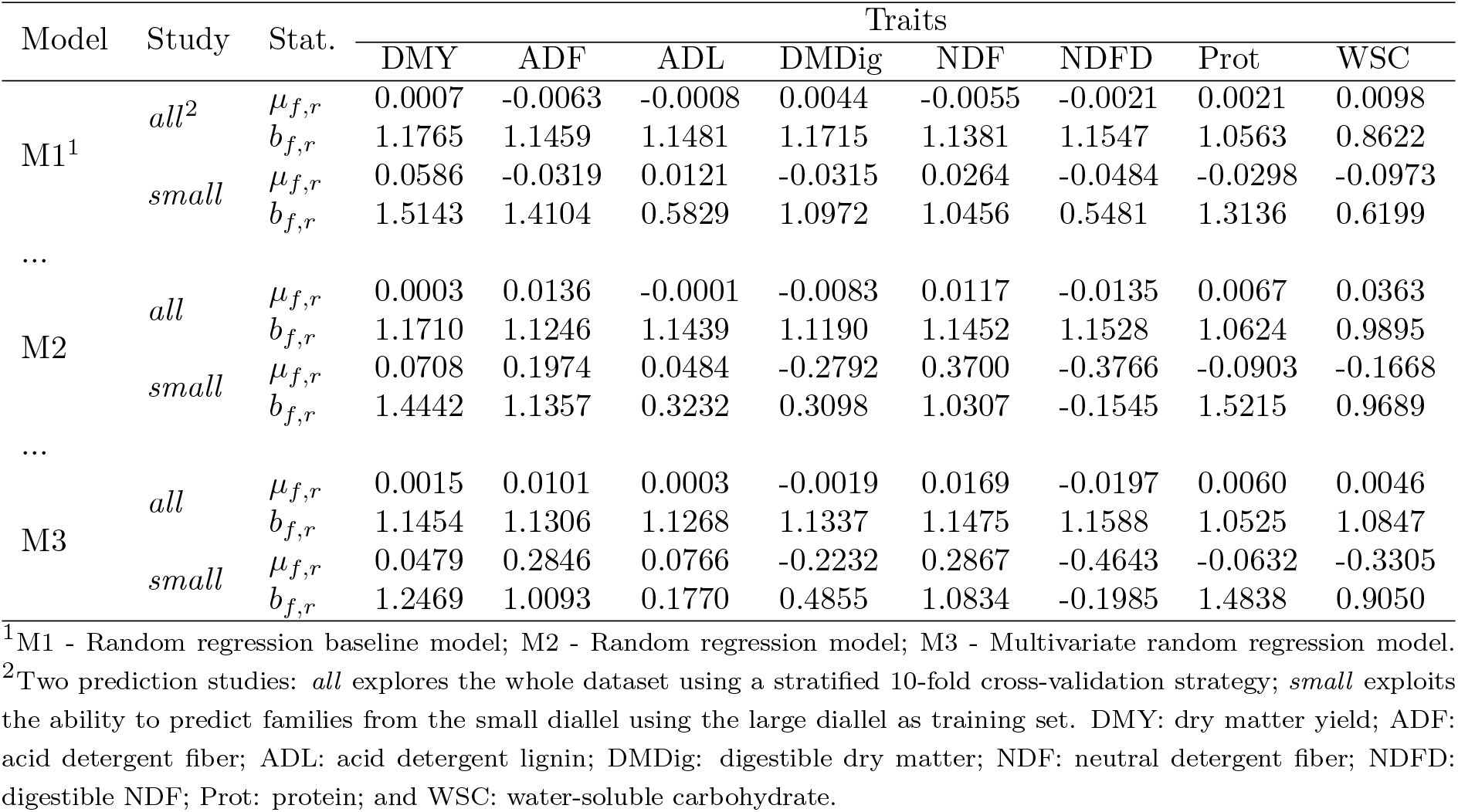
Statistics of prediction bias *µ*_*f,r*_ and dispersion *b*_*f,r*_ estimated as the intercept and slope, respectively, of the linear regression of genomic estimated breeding values (gEBVs) from the full dataset on the gEBVs from the reduced dataset.

| Model | Study | Stat. | Traits |  |  |  |  |  |  |  |
| --- | --- | --- | --- | --- | --- | --- | --- | --- | --- | --- |
|  |  |  | DMY | ADF | ADL | DMDig | NDF | NDFD | Prot | WSC |
| M1<sup>1</sup> | *all*<sup>2</sup> | $\mu_{f,r}$ | 0.0007 | -0.0063 | -0.0008 | 0.0044 | -0.0055 | -0.0021 | 0.0021 | 0.0098 |
| | | $b_{f,r}$ | 1.1765 | 1.1459 | 1.1481 | 1.1715 | 1.1381 | 1.1547 | 1.0563 | 0.8622 |
| | *small* | $\mu_{f,r}$ | 0.0586 | -0.0319 | 0.0121 | -0.0315 | 0.0264 | -0.0484 | -0.0298 | -0.0973 |
| | | $b_{f,r}$ | 1.5143 | 1.4104 | 0.5829 | 1.0972 | 1.0456 | 0.5481 | 1.3136 | 0.6199 |
| ... |  |  |  |  |  |  |  |  |  |  |
| M2 | *all* | $\mu_{f,r}$ | 0.0003 | 0.0136 | -0.0001 | -0.0083 | 0.0117 | -0.0135 | 0.0067 | 0.0363 |
| | | $b_{f,r}$ | 1.1710 | 1.1246 | 1.1439 | 1.1190 | 1.1452 | 1.1528 | 1.0624 | 0.9895 |
| | *small* | $\mu_{f,r}$ | 0.0708 | 0.1974 | 0.0484 | -0.2792 | 0.3700 | -0.3766 | -0.0903 | -0.1668 |
| | | $b_{f,r}$ | 1.4442 | 1.1357 | 0.3232 | 0.3098 | 1.0307 | -0.1545 | 1.5215 | 0.9689 |
| ... |  |  |  |  |  |  |  |  |  |  |
| M3 | *all* | $\mu_{f,r}$ | 0.0015 | 0.0101 | 0.0003 | -0.0019 | 0.0169 | -0.0197 | 0.0060 | 0.0046 |
| | | $b_{f,r}$ | 1.1454 | 1.1306 | 1.1268 | 1.1337 | 1.1475 | 1.1588 | 1.0525 | 1.0847 |
| | *small* | $\mu_{f,r}$ | 0.0479 | 0.2846 | 0.0766 | -0.2232 | 0.2867 | -0.4643 | -0.0632 | -0.3305 |
| | | $b_{f,r}$ | 1.2469 | 1.0093 | 0.1770 | 0.4855 | 1.0834 | -0.1985 | 1.4838 | 0.9050 |

<sup>1</sup>M1 - Random regression baseline model; M2 - Random regression model; M3 - Multivariate random regression model.
<sup>2</sup>Two prediction studies: *all* explores the whole dataset using a stratified 10-fold cross-validation strategy; *small* exploits the ability to predict families from the small diallel using the large diallel as training set. DMY: dry matter yield; ADF: acid detergent fiber; ADL: acid detergent lignin; DMDig: digestible dry matter; NDF: neutral detergent fiber; NDFD: digestible NDF; Prot: protein; and WSC: water-soluble carbohydrate.

